# Exploring the structural basis to develop efficient multi-epitope vaccines displaying interaction with HLA and TAP and TLR3 molecules to prevent NIPAH infection, a global threat to human health

**DOI:** 10.1101/2021.09.17.460735

**Authors:** Sukrit Srivastava, Sonia Verma, Mohit Kamthania, Ajay Kumar Saxena, Kailash C Pandey, Michael Kolbe

## Abstract

**Background:** Nipah virus (NiV) is an emerging zoonotic virus that caused several serious outbreaks in the South Asian region with high mortality rates ranging from 40 to 90% since 2001. NiV infection causes lethal encephalitis and respiratory disease with the symptom of endothelial cell-cell fusion. No specific vaccine has yet been reported against NiV.

**Methodology and Principal Findings:** Recently, the design of some Multi-Epitope Vaccines (MEV) has been proposed but that involves vary limited number of epitopes which limits the potential of vaccine. To address the urgent need for a specific and effective vaccine against NiV infection, in the present study, we have designed two MEVs composed of 33 Cytotoxic T lymphocyte (CTL) epitopes and 38 Helper T lymphocyte (HTL) epitopes. Both the MEVs carry potential B cell linear epitope overlapping regions, B cell discontinuous epitopes as well as IFN-γ inducing epitopes. Hence the designed MEVs carry potential to elicit cell-mediated as well as humoral immune response. Selected CTL and HTL epitopes were validated for their stable molecular interactions with HLA class I and II alleles as well as in case of CTL epitopes, with human transporter associated with antigen processing (TAP). Human β-defensin 2 and β-defensin 3 were used as adjuvants to enhance the immune response of both the MEVs. Molecular dynamics simulation studies of MEVs-TLR3 ectodomain (Toll-Like Receptor 3) complex indicate the stable molecular interaction. Further, the codon optimized cDNA of both the MEVs has shown high expression potential in the mammalian host cell line (Human). Hence for further studies, the designed MEV constructs could be expressed and tried *in-vivo* as potential vaccine candidates against NiV.

**Conclusion:** We conclude that the MEVs designed and in silico validated here could be highly potential vaccine candidate to combat NiV, with greater effectiveness, high specificity and large human population coverage worldwide.

**AUTHOR SUMMARY:** Nipah Virus (NiV) has caused several outbreaks in past two decades calming large number of human lives. Our present work aims to design and *in silico* validate Multi-Epitope Vaccine against NiV. The current approach to design vaccine involves whole virus or full length proteins as vaccine candidates against NiV. These approaches carry chances of raising the unwanted non-neutralizing antibodies which have been found to cause clinical complexities. Recently few Multi-Epitope vaccines have also been proposed, but they have involved limited number of epitopes for vaccine design in result limiting the effectiveness and human population coverage. Here in our MEVs we have involved all the proteins of NiV to design the vaccine. Moreover since we have used in silico validated epitopes we may conclude that the here proposed MEVs would be highly specific, effective and potential vaccine candidate to combat NiV with large human population coverage worldwide.

## INTRODUCTION

Nipah virus (NiV) is an emerging zoonotic virus of the genus Henipavirus of the Paramyxoviridae family [1]. NiV infection causes fatal encephalitis and respiratory disease with a particular symptom of endothelial cell-cell fusion [2]. The first NiV infection to human was first reported in Malaysia in 1998. Later NiV outbreak was reported from Meherpur, Bangladesh in 2001. In the Malaysia NiV infection, the transmission was primarily due human contact with pigs, whereas in later outbreaks of Bangladesh and India the transmission was associated with contaminated date palm sap and human-to-human contact [3]. Bats are identified as the main reservoir for the NiV and they are responsible for the transmission of the infection to both humans and animals [4]. After 2001 NiV outbreak has been reported from different district of Bangladesh almost every year (2003-05, 2007-12). Till March 31, 2012, a total of 209 confirmed cases of NiV infections were reported out of which 161 people died resulting in the mortality rate as high as 77%. After several outbreaks in Bangladesh, in total three NiV outbreaks have also been reported from India. Two of them occurred in the state of West Bengal in 2001 and 2007 [5]. The most recent NiV outbreak was reported from the Kerala state of India during the period of May to June-2018. The Kerala outbreak claimed 17 lives leaving only two survivors out of 19 confirmed cases [6]. Till present, there has been no specific vaccine reported against NiV infection, and the pathogenesis mechanism of NiV to human cells is largely unknown. Hence, an immune-informatics approach investigating the potential of different NiV proteins for vaccine design would be an important and essential step forward for vaccine development.

NiV infection of human cells involves several protein-protein interactions and protein cluster formation on the host cell surface. Essential proteins involved in NiV pathogenesis include C protein, Fusion glycoproteins (F), Glycoproteins (G), Matrix proteins (M), Nucleocapsid protein, Phosphoprotein, Polymerase, V protein and the W protein [7–21]. The C protein regulates the early host pro-inflammatory response as well as the pathogen virulence thus providing a conducive environment for a successful NiV infection [7]. The attachment glycoprotein (G), the fusion protein (F) and the matrix protein (M) together form a cluster on the human cell membrane facilitating virus particle assembly and pathogenesis [8–12]. The G and F proteins of NiV have been shown to be immunogenic by inducing protective immune responses in hamsters [22, 23]. The NiV matrix protein is observed to play a central role in virus particle formation and is essentially required for viral budding from the infected human cells [13–15]. The NiV Polymerase is responsible for the initiation of RNA synthesis, primer extension, and transition to elongation mode and hence the enzyme facilitates viral pathogenesis and survival in host cells [16]. The phosphoprotein and the glycoprotein of NiV are crucially involved in the regulation of viral replication [17, 18] while the V protein of NiV is responsible for the host interferon (IFN) signaling evasion during pathogenesis [19, 20]. Interestingly, the identical N-terminal region of the pathogen’s V and W protein is sufficient to exert the IFN-antagonist activity [21]. Hence, all the nine above mentioned NiV proteins are crucial in different ways for viral pathogenesis and might be important drug and vaccine candidates.

The Nipah virus is a zoonotic RNA virus and it infects human respiratory epithelium cells as well as differentiated neurons (in the brain and spinal cord). Thus, as understood by previous animal model studies, recovery from viral infection and the clearance of viral RNA requires the presence of virus-specific antibodies and interferon gamma (IFN-γ) secretion from T cells [24–26]. Along with the B cell, the T cell also play a critical role in immune response against NiV infection. In recent studies a number of B cell and T cell epitopes from the NIPAH proteome have been reported [27–38]. Further different approaches were proposed for the design of multi epitope vaccines [39, 44]. However, the proposed vaccines utilized a very limited number (6 to 8) of T and B cell epitopes. The use of limited number of epitopes could be challenging for the successful presentation of the exogenous vaccine candidates in view of the proteolytic cleavage by Antigen Presenting Cells (APC).

In the present study we have screened out the most potential Cytotoxic T lymphocyte (CTL) epitopes, Helper T lymphocyte (HTL) and B cell epitopes from the NiV proteome. We have shortlisted and prioritized the most potential and highest scoring 33 CTL, 38 HTL and 16 B cell epitopes. We further evaluated several critical properties (like IC(50), Immunogenicity, Conservancy, Non-toxic etc) to identify the epitopes with the highest immunogeneic potential against NiV. The shortlisted epitopes were utilized for the design of CTL and HTL multi-epitope vaccines against NiV. The designed vaccines were further studied for their stable interaction with innate immunity Toll-Like Receptor 3 (TLR3). The analysis of cDNA of the designed multi-epitope vaccine has predicted to be highly favorable for expression in mammalian cell line. Overall in the present study we have designed and proposed potential multi-epitope vaccines against Nipah virus infection.

## RESULTS & DISCUSSION

### Screening of potential epitopes

#### T cell Epitope Prediction

##### Screening of Cytotoxic T lymphocyte (CTL) Epitope

Cytotoxic T lymphocyte (CTL) epitopes screened and shortlisted according to the highest “Total Score”, low IC(50) (nM) value for epitope-HLA class I allele complexes, and epitopes with the larger number of the HLA class I allele binders. The immunogenicity of the shortlisted CTL epitopes was also determined; the higher immunogenicity score indicates the greater immunogenic potential of the given epitope (Supplementary table S3, S7). In total 33 CD8+ T cell epitopes were finally selected. From this list 10 CTL epitopes (Fusion Protein: FALSNGVLF; Glycoprotein: TVYHCSAVY; Nucleocapsid: YPALALNEF; Phosphoprotein: VSDAKMLSY; Polymerase: YPECNNILF, FPVMGNRIY, AEFFSFFRTF, IPFLFLSAY, ETDDYNGIY, SQNLLVTSY) show a match with previous studies [35–38], indicating consensus of epitope screening by different approaches and methods (Supplementary table S3).

##### Screening of Helper T lymphocyte (HTL) epitopes

The screening of helper T lymphocyte (HTL) epitopes from nine different proteins of NiV was performed on the basis of “Percentile rank”. The smaller the value of percentile rank the higher would be the affinity of the peptide with its respective HLA allele binders. In our initial screening, we got several potential CD4+ T cell epitopes with high scoring. 38 epitopes out of initial screening were shortlisted on the basis that they had highest percentile rank and highest number of HLA class II allele binders (Supplementary table S4, S8).

##### Population Coverage by CTL and HTL epitopes

The population coverage by the shortlisted epitopes was also studied, in particular involving countries of South Asia, East Asia, Northeast Asia and the Middle East. From this study, we concluded that the combined use of all the shortlisted CTL and HTL epitopes would have a cumulative percent of world (countries as listed in Supplementary table S5) population coverage as high as 97.88%.

#### B Cell epitope prediction

##### Sequence-based B Cell epitope prediction

In our initial study, we screened a total of 116 B Cell epitope from nine different NiV proteins, with the epitope length of at least four amino acids utilizing the Bepipred Linear Epitope Prediction method. B cell epitopes predicted by another five different methods based on different physiochemical properties were found to have significant consensus with the epitope amino acid sequences predicted by Bepipred Linear Epitope Prediction. Here, 16 out of the 116 epitopes were shortlisted having a length of 4 to 19 amino acids (Supplementary table S6, Figure 1). One of these 16 B cell epitopes (Matrix Protein: SIPREFMIY) matches with a previous study [39], indicating epitope screening consensus using different approaches and methods (Supplementary table S6).

**Figure 1.**
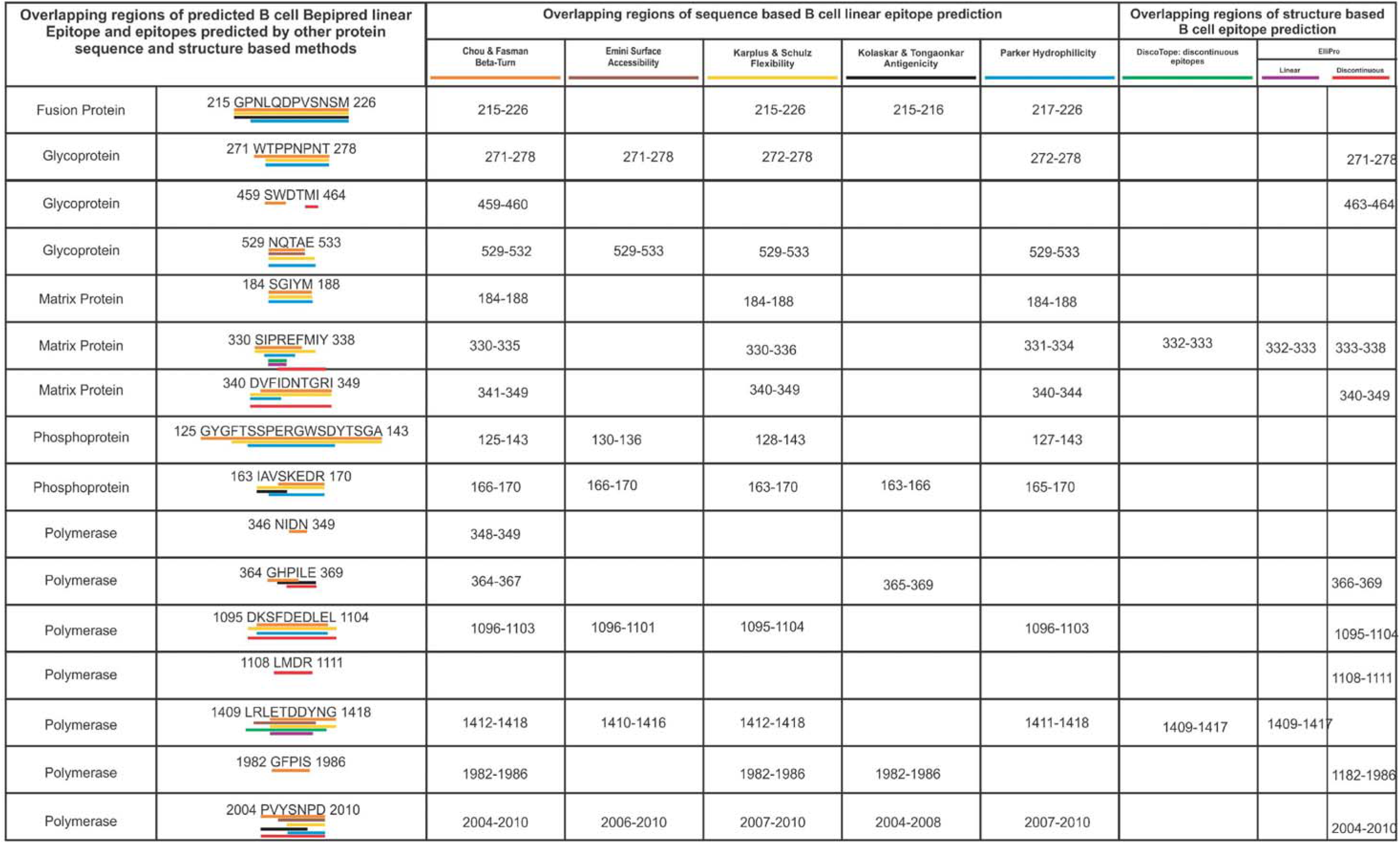
Overlapping regions amongst the linear B cell epitopes predicted by the BepiPred method and other seven different B cell epitopes prediction methods. B cell epitopes predicted by the BepiPred method and other different protein sequence based (Chou.., Emini.., Karplus.., Kolaskar.., and Parker..) and protein structure based (DiscoTope and ElliPro) prediction methods were found to have significant consensus. Consensus overlapping regions of BepiPred epitopes are underlined by the different colour, corresponding to respective prediction method.

##### Structure-based B cell epitope prediction

Structure-based discontinuous and linear epitopes predicted by the DiscoTope 2.0 and the Ellipro methods have shown a significant consensus of overlapping amino acid sequence with the linear epitopes predicted by Bepipred linear epitopes method (Supplementary table S6, Figure 1). This result further confirms that the shortlisted B Cell Bepipred Linear Epitopes have a high chance of causing a strpmg immuno response.

### Characterisation of potential epitopes

#### Epitope conservation and toxicity analysis

Sequence conservation analysis of the shortlisted 33 CTL, 38 HTL and 16 B cell epitopes show that the amino acid sequence conservancy of CTL, HTL and B cell epitopes amongst all retrieved NiV protein sequences is mostly 100%, as shown in Supplementary table S3, S4 and S6. Further, the toxicity analysis of all the shortlisted CTL, HTL and B Cell epitopes was also performed. The ToxinPred study indicated the non-toxic nature of all the shortlisted epitopes (Supplementary table S3, S4, S6).

#### Overlapping residue analysis

Analysis of the amino acid sequence overlap amongst the shortlisted CTL, HTL and B cell epitopes from nine NiV proteins was performed by using the Multiple Sequence Alignment (MSA) analysis tool Clustal Omega. Our analysis showed that several epitopes of CTL, HTL and B cell have overlapping amino acid sequence. Besides a few shorter identical sequences the majority of the CTL, HTL and B cell epitopes have several amino acids overlapping as shown in Figure 2.

**Figure 2.**
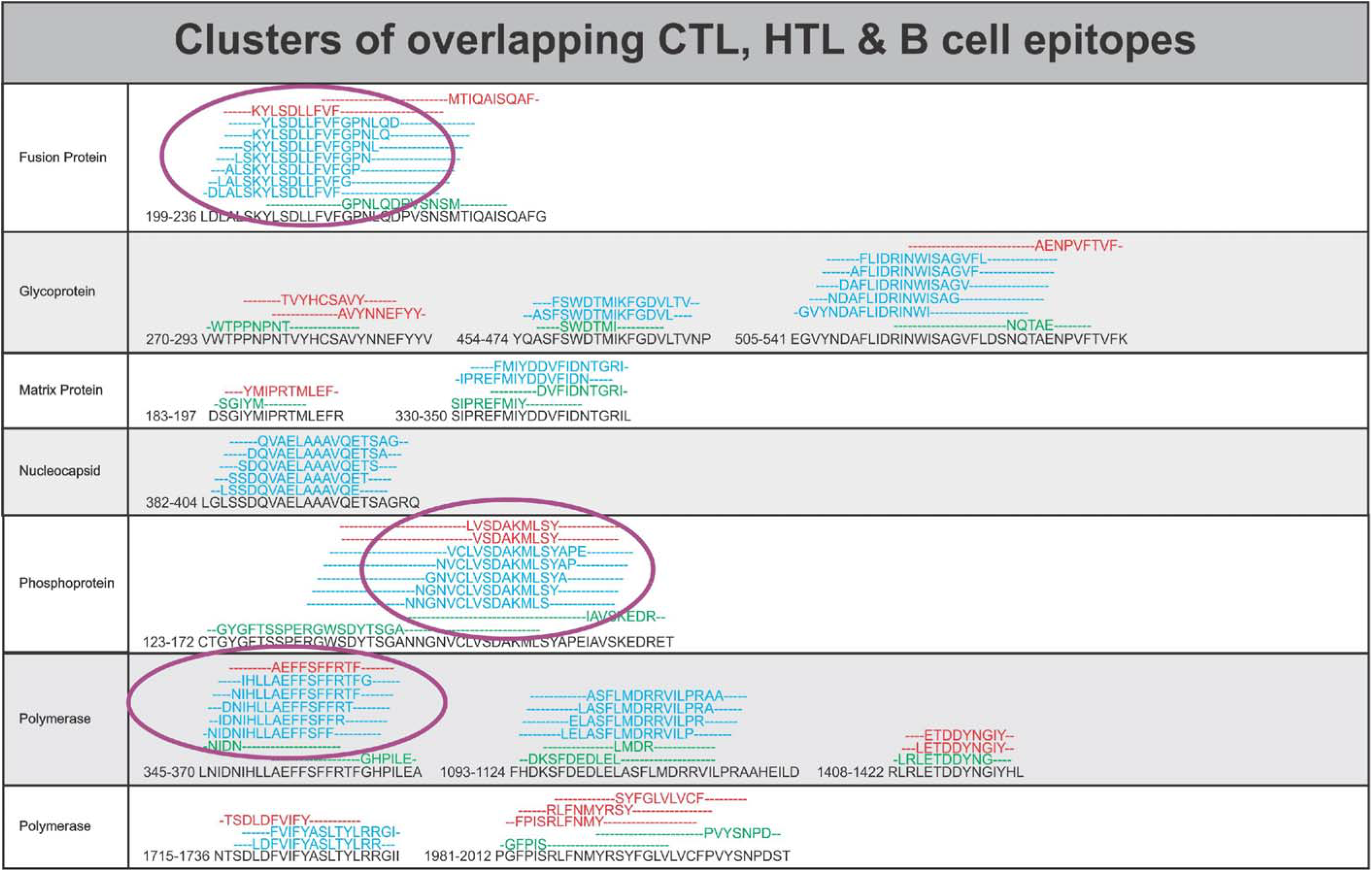
Overlapping CTL, HTL and B cell epitopes. Multiple sequence alignment performed by Clustal Omega at EBI to identify the consensus overlapping regions of CTL (red), HTL (blue) and B cell epitopes (green) amongst shortlisted epitopes. Epitopes with overlapping region amongst all the three types of epitopes (CTL, HTL and B Cell epitopes), epitopes with full sequence overlap and epitopes with the highest number of HLA allele binders were chosen for further studies (encircled).

#### Epitope selected for molecular interaction study with HLA allele and TAP transporter

Amongst all the shortlisted epitope peptides seven CTL and seventeen HTL epitope peptides have partial or full overlapping sequences or have the highest number of HLA allele binders were shortlisted for further studies (Figure 2, Supplementary table S3 & S4).

### Molecular interaction analysis of selected epitopes with HLA allele and TAP transporter

#### Molecular interaction analysis of selected CTL and HTL epitopes with HLA alleles

The molecular docking study of the shortlisted CTL and HTL epitopes with their respective HLA class I and II allele binders was performed using AutoDock Vina. Docking studies revealed for all epitopes significant molecular interactions with their HLA allele binders having low binding energies and multiple hydrogen bonds formed (Figure 3A & 4B). The stability of the obtained docking complexes was further tested by molecular dynamics (MD) simulation studies. MD simulations were performed over a time interval of 0.5-1 ns at the invariable temperature of ∼ 300 K and at invariable pressure of ∼ 1 bar. All the complexes showed reasonably invariant root mean square deviation (RMSD) value (between ∼ 0.2 to 0.4 nm) indicating the stable nature of the tested epitope-HLA allele complexes (Figure 4A & 5B). Moreover, the reasonably invariant Rg (radius of gyration) of the complexes, throughout the MD simulation (Supplementary figure S2), and the root mean square fluctuation (RMSF) for all the atoms of the complexes (Supplementary figure S3) again indicate the stable nature of the epitopes and HLA allele complexes. Furthermore, the B-factor analysis of all the epitope-HLA allele complexes indicated most of the complex regions to be stable (blue) with a very small region being acceptably fluctuating (yellow and orange) (VIBGYOR color presentation) Supplementary figure S4.

**Figure 3.**
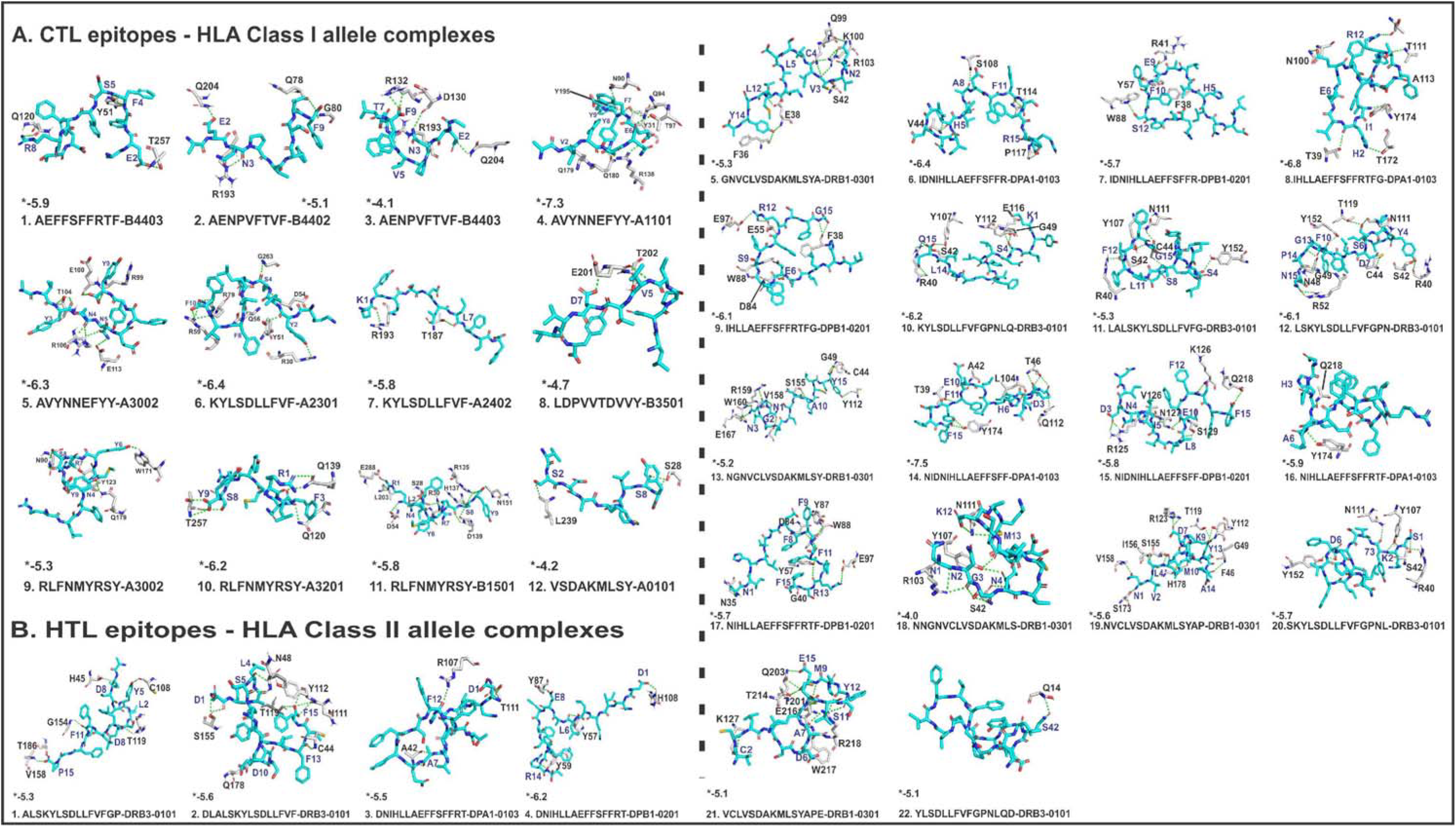
**(A). Molecular Docking analysis of CTL epitopes and HLA alleles.** Molecular docking of selected CTL epitopes (cyan sticks) with their respective HLA class I allele binders (gray sticks). The study shows the docked complexes to have significantly negative binding energy along with hydrogen bonds (green dots) formation in the complex interface. **(B) Molecular Docking analysis of HTL epitopes and HLA alleles.** Molecular docking of selected HTL epitopes (cyan sticks) with their respective HLA class II allele binders (gray sticks). The study shows the docked complexes to have significantly negative binding energy along with hydrogen bonds (green dots) formation in the complex interface. (*) Indicates binding energy, shown in kcal/mol.

**Figure 4.**
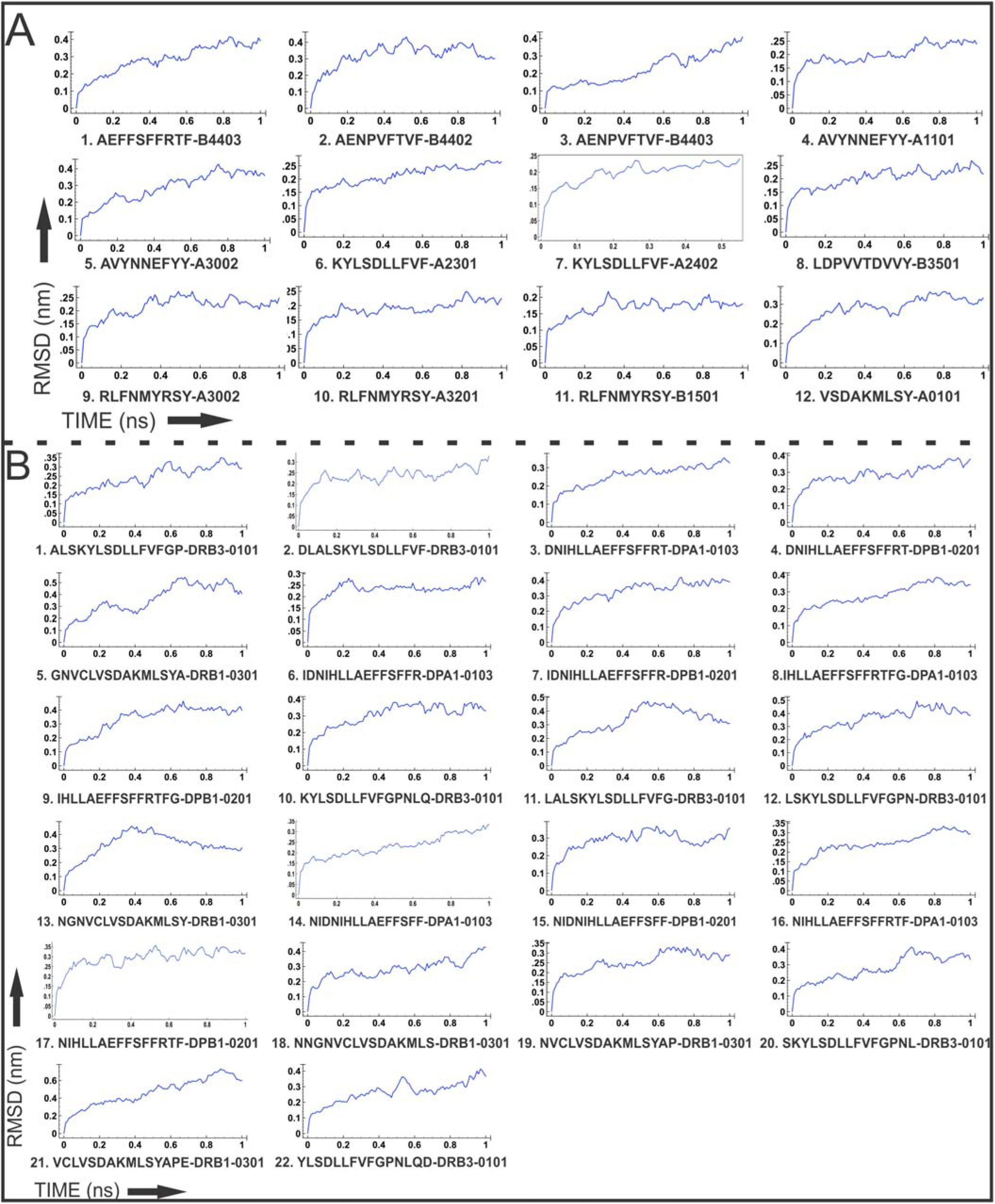
**(A). Molecular Dynamics simulation analysis of CTL epitopes and HLA allele complexes.** Molecular Dynamics simulation study reveals a stable nature of the CTL-HLA allele complexes throughout 0.5-1 ns time window with reasonably invariable RMSD. **(B) Molecular Dynamics simulation analysis of CTL epitopes and HLA allele complexes.** Molecular Dynamics simulation study reveals a stable nature of the HTL-HLA allele complexes throughout 0.5-1 ns time window with reasonably invariable RMSD.

**Figure 5.**
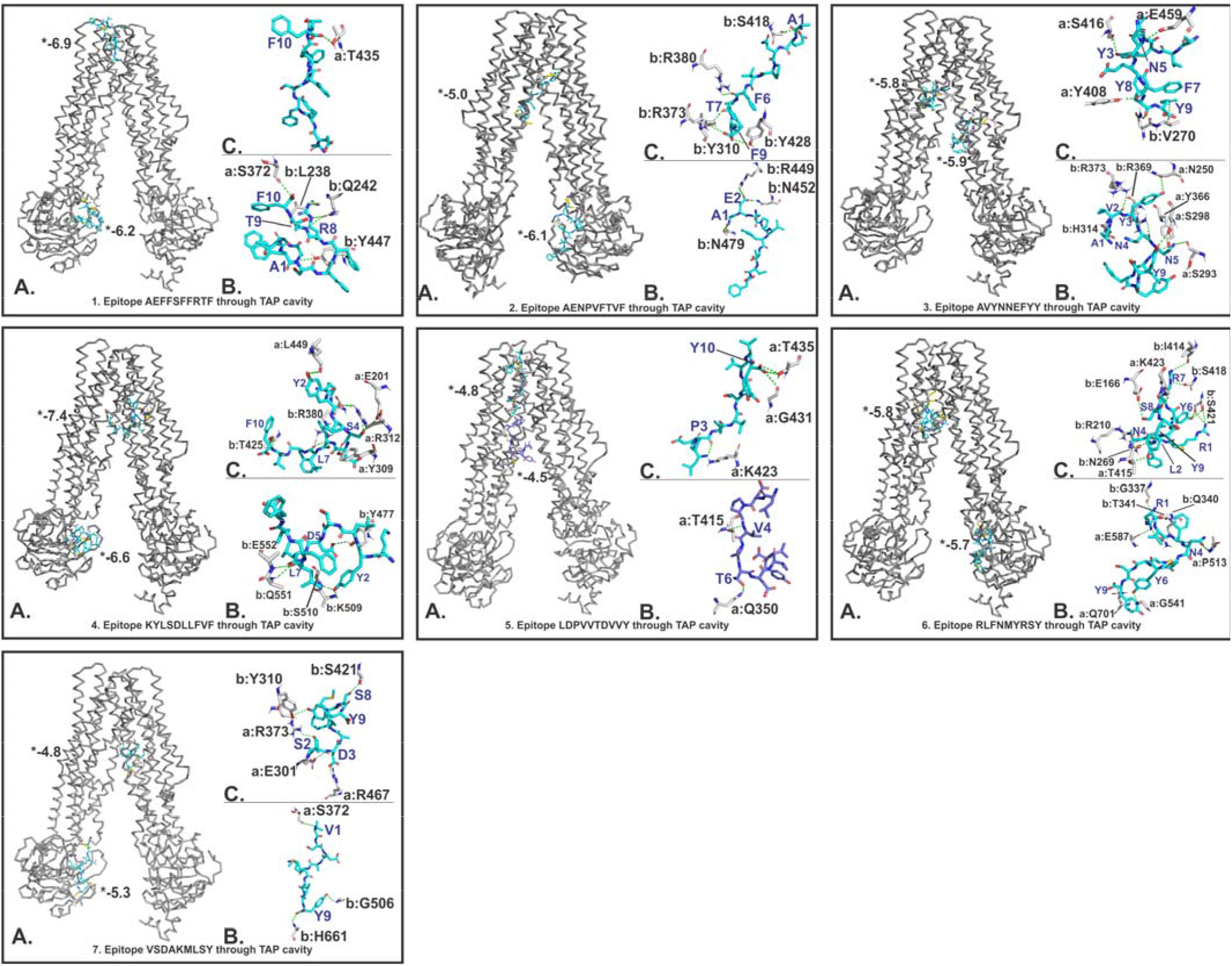
Molecular docking analysis of CTL epitopes within the TAP transporter cavity. Molecular interaction of CTL epitopes (cyan sticks) within the TAP cavity (gray ribbon/sticks) is shown in detail. For every panel of epitope-TAP complex, **(A)** shows the binding of epitope at two different sites within TAP cavity, **(B)** and **(C)** show detailed molecular interaction between epitopes and TAP cavity; (a, b) show chain A and B of TAP transporter. H-bonds are shown in yellow dots. (*) Indicates binding energy, shown in kcal/mol.

#### Molecular interaction analysis of selected CTL epitopes with TAP transporter

The molecular docking interaction analysis of the chosen CTL epitopes with the TAP transporter cavity showed a strong molecular interaction with low binding energy and several hydrogen bonds formed at different sites of the TAP transporter cavity. Two sites of interaction were of particular interest, one located near the cytoplasmic end and the other in the vicinity of the ER lumen (Figure 5). Our study confirms the transportation feasibility of the chosen CTL epitopes from the cytoplasm into the ER lumen which is essential for the representation of peptides by the HLA allele molecules on the surface of antigen presenting cells.

### Design, characterisation and molecular interaction analysis of Multi-Epitope Vaccines with immunological receptor

#### Design of Multi-Epitope Vaccines

All the screened CTL and HTL epitopes were utilized to design CTL and HTL Multi-Epitope vaccines. Short peptides EAAAK and GGGGS were used as rigid and flexible linkers respectively (Figure 6). The GGGGS linker provides proper conformational flexibility to the vaccine tertiary structure and hence facilitates stable conformation to the vaccine. The EAAAK linker facilitates in domain formation and hence facilitates the vaccine to obtain its final stable structure [45–54]. The human beta-defensin 2 (hBD-2) (PDB ID: 1FD3, Sequence: GIGDPVTCLKSGAICHPVFCPRRYKQIGTCGLPGTKCCKKP) and the human beta-defensin 3 (hBD-3) (PDB ID: 1KJ6, Sequence: GIINTLQKYYCRVRGGRCAVLSCLPKEEQIGKCSTRGRKCCRRKK) were used as adjuvants in the design of both the MEVs at N and C terminals respectively [45-54,]. Human Beta-defensins (hBD) have an important role in the chemotactic activity memory T cells, immature dendritic cells and monocytes and they are involved in the degranulation of mast cells. Due to the important role of hBDs in immune response enhancement, hBDs have been chosen and utilized as adjuvants for the MEV designs.

**Figure 6.**
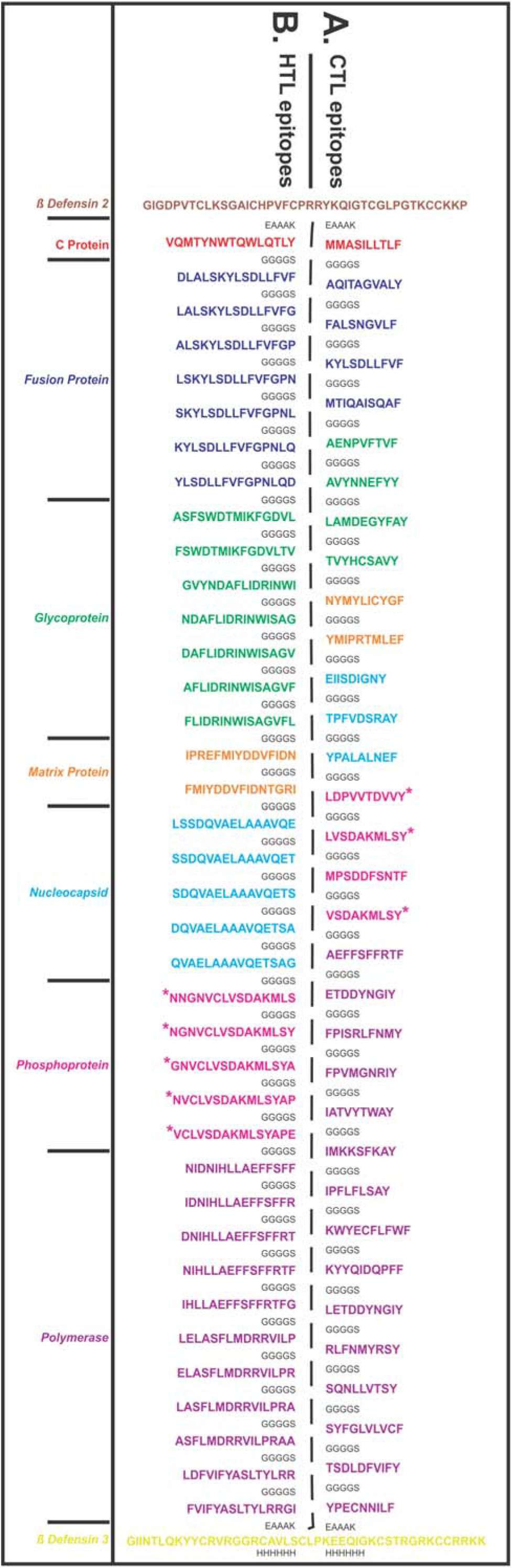
Design of Multi-Epitope Vaccine (MEVs). **(A)** CTL and **(B)** HTL epitopes were linked by the short peptide linker ‘GGGGS’. Human β Defensin 2 and β Defensin 3 were used as an adjuvant at the N and C terminals respectively. The short peptide EAAAK was used to link the β Defensin 2 and β Defensin 3. Epitopes from different proteins were coloured in different colours. C terminal 6xHis is designed as His tag. * Epitopes common to Phosphoprotein, V Protein and W protein.

### Characterisation and molecular interaction analysis of designed Multi-Epitope Vaccines with immunological receptor

#### Characterisation of designed Multi-Epitope Vaccines

##### Interferon-gamma inducing epitope prediction

Interferon-gamma (IFN-γ) inducing epitopes are involved in both the adaptive and the innate immune response. The IFN-γ inducing 15-mer peptide epitopes were screened from the CTL and HTL MEVs by utilizing the IFNepitope server. A total of 33 CTL MEV and 43 HTL MEV INF-γ inducing POSITIVE epitopes with a score of 1 or more than 1 were shortlisted (Supplementary table S9, Figure 7D & 7I).

**Figure 7.**
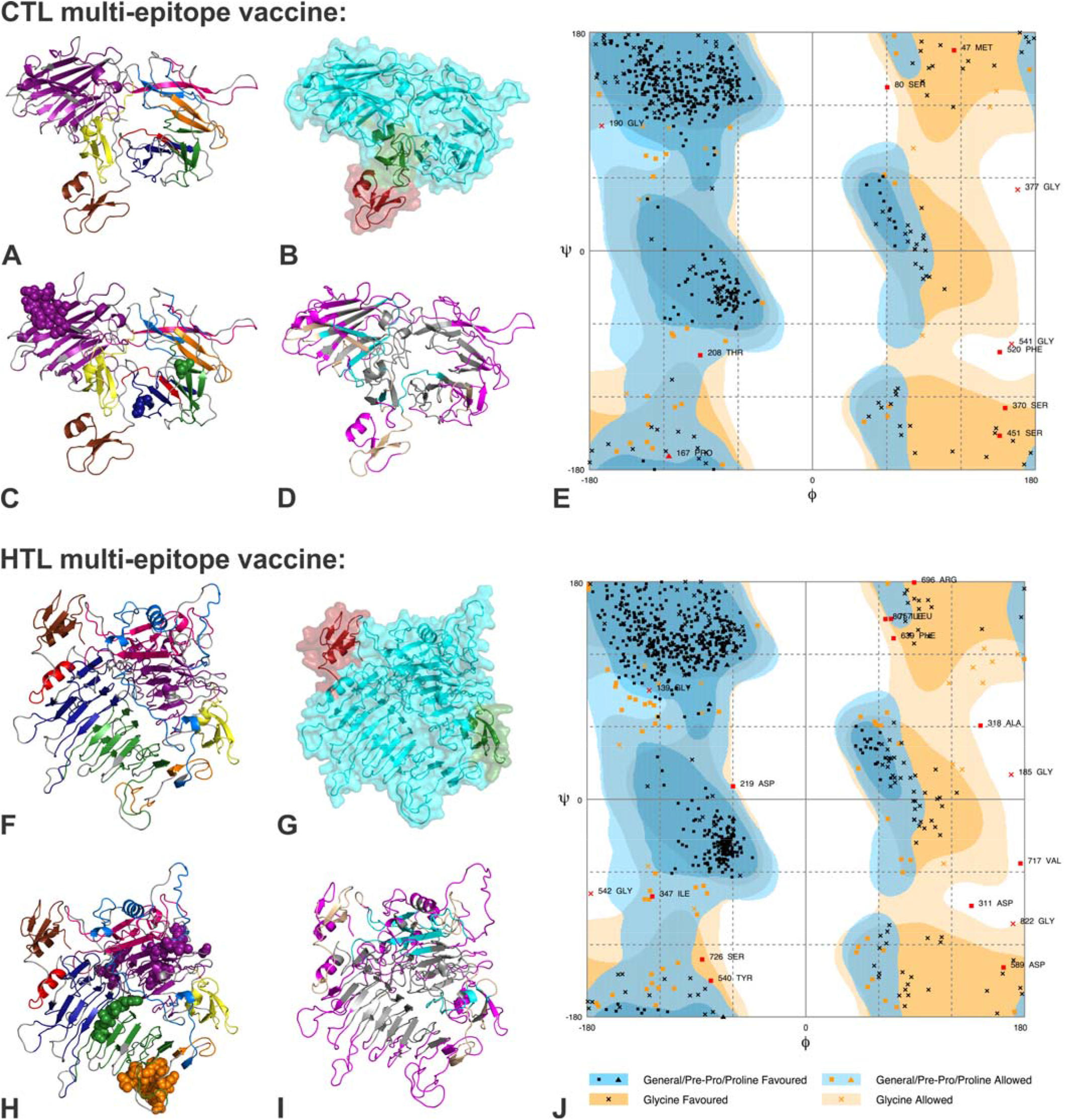
Tertiary structure modelling of CTL and HTL Multi-Epitope Vaccines. **(A) & (F)**: Tertiary structural models of CTL and HTL MEVs showing epitopes in different colours corresponds to as in Fig. 1. **(B) & (G)**: Show the different domains of CTL and HTL MEVs. **(C) & (H)**: The overlapping linear B cell epitope region present in CTL and HTL MEVs, shown by spheres. **(D) & (I):** From the CTL and HTL MEVs, the INF-γ inducing epitopes are shown in cyan, discontinuous B Cell epitopes are shown in magenta and the region common amongst INF-γ and discontinuous B Cell epitopes are shown in wheat colour. **(E) & (J)**: RAMPAGE analysis of the refined CTL and HTL MEV models.

##### MEVs allergenicity and antigenicity prediction

Both the CTL and HTL MEVs were analyzed to be NON-ALLERGEN by the AlgPred analysis scoring −0.612 and −0.935 respectively while default threshold value being −0.4. CTL and HTL MEVs were also analyzed by VaxiJen to be probable ANTIGENS with the prediction score of 0.4447 and 0.4836 respectively, while the default threshold value for viral proteins being 0.4. Hence, with the mentioned analysis tools both the CTL and HTL MEVs are predicted to be non-allergic as well as antigenic in nature.

##### Physicochemical property analysis of designed MEVs

ProtParam calculations were performed for both the CTL and HTL MEVs to analyse their physiochemical properties. The CTL MEV is composed of 576 amino acids, has a molecular weight of 58.57 kDa and a theoretical pI of 8.19. The expected half-life of the CTL MEV in *E.coli*, yeast and mammalian reticulocytes were predicted with 10 h, 20 min, and 30 h respectively; the aliphatic index of CTL MEV was found to be 58.42, and grand average of hydropathicity (GRAVY) of CTL MEV was found to be −0.010, both indicating globular and hydrophilic nature of the CTL MEV. The instability index score of the CTL MEV was 48.03 indicating its stable folding under native conditions. Further, the ProtParam analysis of the HTL MEV calculated for the 857 amino acids, a molecular weight of 87.62 kDa and a theoretical pI of 5.99. The expected half-life of HTL MEV in *E.coli*, yeast and mammalian reticulocytes was predicted to be 10 h, 20 min and 30 h, respectively. The aliphatic index of HTL MEV was calculated as 82.99, and the grand average of hydropathicity (GRAVY) of the HTL MEV was found to be 0.188, both indicating that HTL MEV has a globular and hydrophilic nature. The instability index of the HTL MEV was 45.66 indicating its stable nature.

Overall, the physiochemical properties of both proteins indicate that both the proteins might be natively expressed as stable proteins and purified with standardized purifications methods and protocols.

##### Tertiary structure modelling and refinement of MEVs

Tertiary structural models were generated for both the CTL and HTL MEVs by utilizing the RaptorX modelling tool (Figure 7). The model obtained for CTL MEV has 6% helix, 27% β-sheet, 66% coil content and structural elements are 23% exposed, 39% medium and 36% buried. The structural model of CTL MEV has three domains ranging from amino acid 1 to 46 (1^st^ domain, template-1fd3:A), 47 to 520 (2^nd^ domain, templates-1yrzA, 1y7bA, 5jozA, 3zxjA, 5z5dA) and 521 to 576 (3^rd^ domain, template-1kj6:A) (Figure 7B). Similarly, the 3D model calculated with RaptorX for the HTL MEV has 21% helix, 22% β-sheet, 56% coil content with 24% of the amino acids exposed, 39% medium and 36% buried. The structural model of HTL MEV has three domains ranging from amino acid 1 to 46 (1^st^ domain, template-1fd3:A), 47 to 801 (2^nd^ domain, templates-5m5zA, 3eqnA), 802 to 857 (3^rd^ domain, template-1kj6:A) (Figure 7G). The P-Value for the best template based CTL and HTL MEV homology models were 2.79e-04 and 5.99e-03 respectively. Good quality, mostly alpha proteins have a P-value of less than 10^-3^ and that of mostly beta proteins has a P-value of less than 10^-4^. Hence, both the homology models of CTL and HTL MEVs are predicted to be of good quality. Since for the CTL and HTL MEV design, the CTL and HTL epitopes used also show overlapping common regions with the linear B cell epitopes (Figure 1), both the generated CTL and HTL MEV models also carry the overlapping regions of linear B Cell epitopes (Figure 7C, 7H).

The generated CTL and HTL 3D models were further refined using ModRefiner to fix structural gaps followed by GalaxyRefine refinement. Refinement with ModRefiner showed a TM-score of 0.9703 and 0.8934 for the CTL and HTL models, respectively. Since both values are close to 1 this indicates that the initial and the refined models were structurally similar. For the CTL MEV model refinement, the sores of model 1 for different parameters were, Rama favored: 90.8%, GDT-HA: 0.9596, RMSD: 0.385, MolProbity: 2.673, Clash score: 29.9, and Poor rotamers: 1.8. Likewise for HTL MEV model refinement, the sores of model 1 for different parameters were, Rama favoured: 88.5%, GDT-HA: 0.9463, RMSD: 0.419, MolProbity: 2.811, Clash score: 38.4, and Poor rotamers: 1.6. Here, MolProbity shows the log-weighted combination of the clash score, percentage Ramachandran not favoured and the percentage bad side-chain rotamers. After refinement, all the mentioned parameters were significantly improved in comparison to the initial CTL and HTL MEV models (Supplementary table S10).

##### Validation of CTL and HTL MEVs refined models

Both the CTL and HTL model were analysed with the RAMPAGE analysis tool after refinement. The refined CTL MEV model has 91.5% residues in favored regions and the refined HTL MEV model has 89.5% of residues in favored region. This indicates both the CTL and HTL MEVs have gained an acceptable conformation after refinement (Figure 7E, 7J).

##### Discontinuous B-cell epitope prediction from MEVs

Discontinuous B-cell epitopes were further predicted from the final refined 3D models of CTL and HTL MEVs utilizing the ElliPro tool on IEDB server. The screening revealed that the CTL MEV carries 3 and the HTL MEV has 2 potential discontinuous epitopes. The PI (Protrusion Index) score of the CTL MEV discontinuous B cell epitopes ranges from 0.682 to 0.747 and that of HTL MEV it ranges from 0.687 to 0.745 (Supplementary table S11, Figure 7D & 7I). The higher PI score indicates a greater potential of the discontinuous B cell epitope.

#### Molecular interaction analysis of MEVs with TLR-3

The refined models of CTL and HTL MEVs were further studied for their molecular interaction with the ectodomain (ECD) of human TLR-3. Therefore, molecular docking of CTL and HTL MEVs model with the TLR-3 crystal structure model (PDB ID: 2A0Z) was performed utilizing the PatchDock tool. Generated docking conformation with highest geometric shape complementarity score of 22382 and 18264 for CTL and HTL MEVs, respectively were selected for further studies. The highest docking score predicted with the PatchDock tool indicates the best geometric shape complementarity fitting conformation of MEVs and the TLR-3 receptor. Both, the CTL and HTL MEVs were fitting into the ectodomain region of TLR-3 after docking (Figue 8A & 8E). The CTL and HTL MEVs have shown to form multiple potential hydrogen bonds within the ectodomain cavity region of TLR-3. Further, the molecular dynamics simulation study was also performed for the docked complexes of both the MEVs and TLR-3. In MD simulations both the complexes have shown reasonably stable effective RMSD value between ∼ 0.2 to 4 Å for a given time window of 100 ns at invariable pressure (∼ 1 bar) and temperature (∼ 300 K) (Figure 8B & 8F). The effective RMSD is well within tange of Hydrogen bond formation, moreover the range is towards 0.2 Å for most of duration. The reasonably invariant radius of gyration (Rg) (∼ 0.7 to 2 Å) of both MEVs-TLR-3 complexes (Figure 8C & 8G), and RMS fluctuation (RMSF) (∼ 2 to 3 Å) for all the atoms in both the complexes (Figure 8D & 8H) indicate that the MEVs-TLR3 complexes are stable. The B-factor analysis of MEVs-TLR3 complexes was also performed. The B-factor indicates the displacement of the atomic positions from an average (mean) value i.e. the more flexible an atom is the larger the displacement from the mean position will be (mean-squares displacement) (Figure 8A and 8E). The areas with high B-factors are colored red (hot), while low B-factors are colored blue (cold) (VIBGYOR presentation). The B-factor of most of the regions of MEVs-TLR3 complexes indicates the stable nature of the complexes while a very small region is found to be fluctuating. The results suggest a stable complex formation tendency for both the CTL and HTL MEVs with the ectodomain of the human TLR-3 receptor.

**Figure 8.**
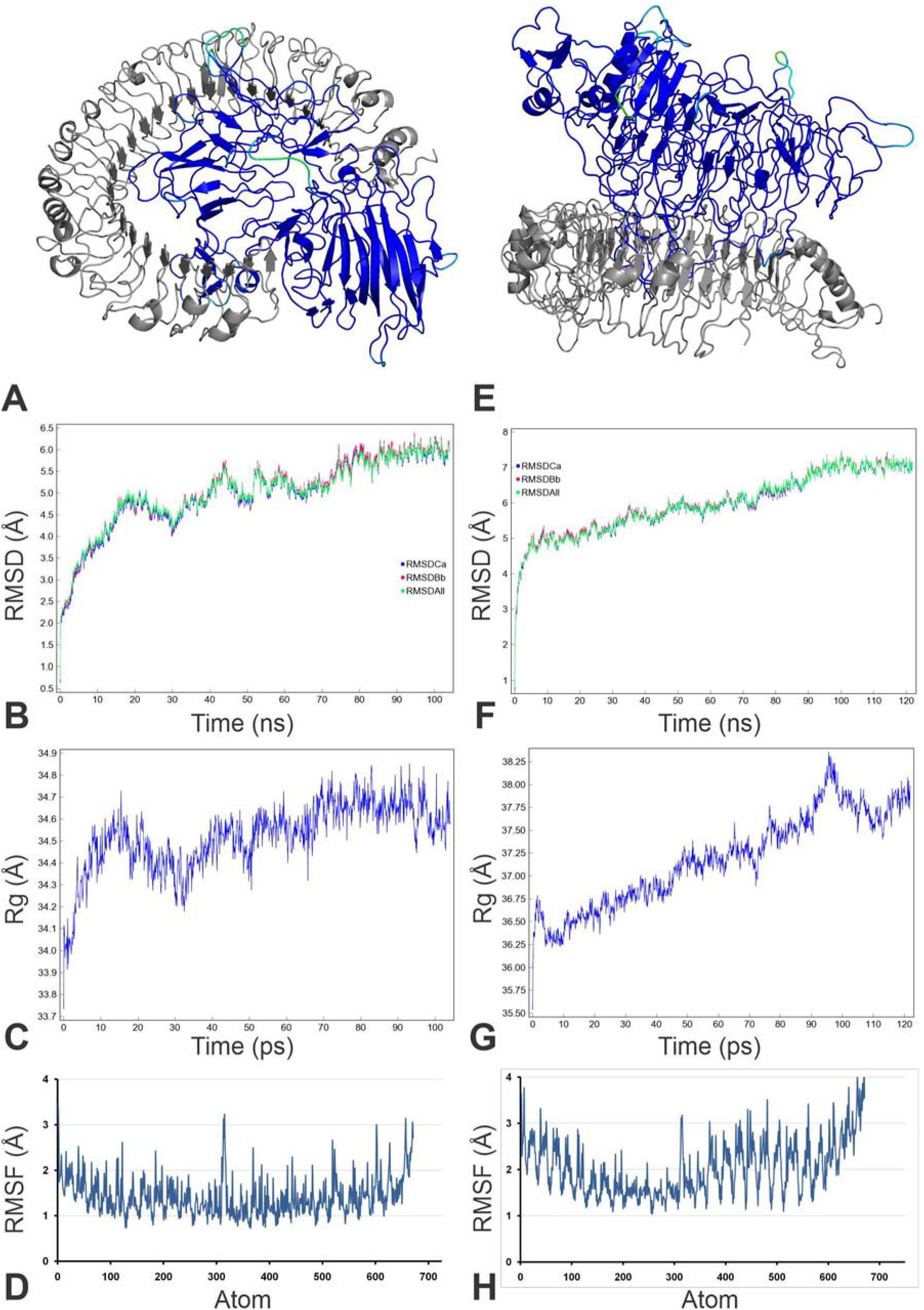
Molecular Docking and dynamics simulation study of CTL and HTL MEVs with TLR-3. **(A)** CTL and **(E)** HTL MEVs (VIBGYOR) docked complex with TLR-3 (gray). Both the complexes are forming several hydrogen bonds in the MEV and TLR-3 interface, as shown by green dots. B-Factor of the docked MEVs is shown by a rainbow (VIBGYOR) presentation. The regions in blue being indicated stable and the region in red indicate unstable. In the above complexes, most of the region of docked MEVs is in blue and with the very small region is green, yellow or orange, hence the complexes are predicted to be very stable. **(B)** and **(F),** RMSD as generated by the molecular Dynamics simulation study of CTL, HTL MEVs and TLR-3 complexes. **(C)** & **(G)** Rg (radius of gyration) across the time window of 100 nanosecond. **(D)** & **(H)**, RMS fluctuation for all the atoms of the CTL, HTL MEVs and TLR-3 complexes.

#### *In-silico* analysis of MEVs for cloning and expression

##### Analysis of cDNA of both the MEVs for cloning and expression in mammalian cell line

Complementary DNA codon optimized for CTL and HTL expression in mammalian host cell lines (Human) was generated with the Java Codon Adaptation Tool. The generated optimized cDNA’s for both the MEVs were also analysed by utilizing the GenScript Rare Codon Analysis Tool revealing aGC content of optimized CTL-MEV cDNA of 69.79% and a CAI (Codon Adaptation Index) score of 1.00 with 0% tandem rare codons. Likewise, the GC content of the optimized HTL-MEV cDNA was 70.69%, CAI score was 1.00 with 0% tandem rare codons. Since for higher possibility for cDNA expression in human expression system, the GC content of a cDNA should be within the range of 30% to 70%, the CAI score should be between 0.8-1.0, and the tandem rare codon frequency that indicates the presence of low-frequency codons, should be <30%, the cDNA constructs of CTL and HTL MPVs are expected to have high potential for expression in human expression system. The tandem rare codons may hinder the proper expression of the cDNA or even interrupt the translational machinery of the chosen expression system. According to the GenScript Rare Codon Analysis the cDNA of both the MEVs satisfy all the mentioned parameters and are predicted to have high expression in the mammalian host cell line (Human).

## CONCLUSION

In the present study, we have designed and validated two multi-epitope vaccines derived from CTL and HTL epitopes. The selected peptides show significant sequence overlap with screened linear B cell epitopes. Both, the CTL and HTL MEVs tertiary models carry potential discontinuous B cell epitopes and INF-γ epitopes. Consequently, the designed MEVs might have the potential to elicit profound humoral and cellular immune responses. Human β-Defensin 2 and 3 fused to the N and C terminal ends of both the MEVs serving as adjuvants to enhance the immune response. The identified epitopes for the CTL and HTL MEVs were also validated by molecular docking and MD simulation studies to test the interaction with their respective HLA allele binders. Molecular interaction of the selected CTL epitopes within the TAP transporter cavity was also evaluated indicating a favorable transport of epitopes from cytoplasm to lumen of Endoplasmic Reticulum for further presentation on cell surface by Golgi bodies. Our analysis of the shortlisted CTL and HTL epitopes combined revealed coverage of 97.88% world human population. The molecular interaction analysis of both the CTL and HTL MEVs with the immunoreceptor TLR3 showed structural fit of the MEVs into the ectodomain of TLR3 receptor cavity and MD simulations indicate very stable complexes formed. Since both the CTL and HTL MEVs carry CTL, HTL as well as discontinuous B cell epitopes, the combined administration of both the MEVs, is predicted to elicit both the humoral as well as cell-mediated immune response. The cDNA for both the MEVs were designed considering codon-bias for the expression in mammalian host cell lines (Human). The cDNAs were optimized with respect to their GC content and zero tandem rare codons to gain high expression in mammalian host cell lines (Human). In future experimental studies the designed CTL and HTL MEVs should be cloned, expressed and validated *in-vivo* and in animal trails as potential vaccine candidates against NiV infection.

## METHODOLOGY

In the present study, we have designed two multi-epitope vaccines (MEVs) composed of thoroughly screened most potential Cytotoxic T lymphocyte (CTL) epitopes and Helper T lymphocyte (HTL) epitopes derived from the nine NiV proteins (glycoprotein (gi-253559848); C protein (gi-1859635642); fusion protein (gi-13559813); matrix protein (gi-13559811); nucleoprotein (gi-1679387250); phosphoprotein (gi-1802790259); V protein (gi-1802790260); RNA polymerase (gi-15487370); W protein (gi-374256971). CTL and HTL epitopes would be the most potential vaccine candidates since they are responsible for cell-mediated immune response by their presentation on the surface of antigen presenting cells (APCs) by their respective Class I and II human leukocyte antigen (HLA) allele binders. Both the CTL and HTL epitopes chosen for MEV design also carry overlapping regions of linear B cell epitopes. Moreover, both the MEVs also carry potential discontinuous B cell epitopes as well as IFN-γ inducing epitopes in their tertiary structure model. Hence, both the designed MEVs carry potential to elicit cell-mediated as well as humoral immune response. Furthermore, both the MEVs were designed with the vaccine adjuvant human β Defensin 2 and human β Defensin 3 at their N and C termini to enhance the immune response [45–46]. β-defensins have considerable immunological adjuvant activity in general and, in particular the β-defensins 2 & 3 have been shown in previous studies to generate potent humoral immune responses when fused with B-cell lymphoma epitopes in mouse model [47–50]. Since, the pro-inflammatory mediators enhance the expression of β-defensins 2 & 3 in airway epithelial cells we chose β-defensins in our study. The selected CTL and HTL epitopes were validated for their stable molecular interactions with their respective HLA alleles binders; for CTL epitopes, their molecular interaction with human Transporter associated with antigen processing (TAP) was also analyzed [55, 56]. This analysis validated the CTL epitopes that get transported through the TAP cavity for their presentation on cell surface or not. The human TAP selectively transports cytosolic peptides into the lumen of the endoplasmic reticulum in an ATP-dependent manner [57]. Tertiary structure models of both the MEVs were generated, refined, and further docked with the human Toll-Like Receptor 3 (TLR3), which is an essential immuno-receptor in this pathway [58, 59]. The nuclear localization of Nipah virus W protein inhibits the signaling pathway of TLR3 upon pathogenesis to suppress the TLR3 induced activation of the IFN response to eventually prevent relay of the warning signals to uninfected cells. TLR3 is preferentially expressed by human astrocytes of the central nervous system (CNS) upon infection. The NiV infection involves invasion of the neurons of CNS and hence causes infection in CNS. These studies indicate the importance of the TLR3 responses in immune response and hence TLR3 has been chosen to be studied for its stable binding with the designed multi-epitope vaccines [60–63]. The complexes of CTL and HTL MEVs formed with the human TLR3 were further analysed for their stable molecular interaction by a molecular dynamics simulation study. The cDNA of the designed MEVs were generated and analysed for their high expression tendency in the mammalian host cell line (Human). Overall, from the present *in-silico* study, we may put forward the design of two MEVs, which qualify all the significant criterions for being a potential vaccine candidate against NiV infections. The corresponding workflow is shown in Supplementary figure S1.

### NiV proteins selected for potential epitope screening

In the present study, nine NiV proteins were used for epitope screening. They include C protein, Fusion glycoprotein (F), Glycoprotein (G), Matrix proteins (M), Nucleocapsid protein, Phosphoprotein, Polymerase, V protein and the W protein. The full-length protein sequences of NiV proteins were retrieved from the NCBI database (National Center for Biotechnology Information (https://www.ncbi.nlm.nih.gov/protein). A total of 161 protein sequences available at NCBI, belonging to different strains and origins of NiV, were retrieved. For structural based epitope screenings available tertiary structures of NiV proteins were retrieved from Protein Data Bank (PDB) (http://www.rcsb.org/pdb/home/home.do). For the NiV proteins with no structure available, homology models were generated by Swiss-model, (http://swissmodel.expasy.org/) [64] (Supplementary table S1).

### Screening of Potential Epitopes

#### T cell Epitope Prediction

##### Screening of Cytotoxic T lymphocyte (CTL) Epitope

The screening of Cytotoxic T lymphocyte epitopes was performed by the IEDB (Immune Epitope Database) tool “Proteasomal cleavage/TAP transport/MHC class I combined predictor” (http://tools.iedb.org/processing/) [65–67]. Proteasome cleavage score depends on the total amount of cleavage site in the protein. TAP score estimates an effective log -(IC50) values (half maximal inhibitory concentration (IC50)) for binding to TAP of a peptide or its N-terminal prolonged precursors. The MHC binding prediction score is −log(IC50) values for binding to MHC of a peptide [68]. The tool gives a “Total Score” which is a combined score of the proteasome, MHC, TAP (N-terminal interaction), and processing analysis scores. The total score is generated by using the combination of six different methods viz. Consensus, NN-align, SMM-align, Combinatorial library, Sturniolo and NetMHCIIpan [65–68]. The IC(50) (nM) value for each epitope and MHC allele binding pairs were also obtained by this IEDB tool. Epitopes having high, intermediate, and least affinity of binding to their HLA allele binders have IC50 values < 50 nM, < 500 nM and < 5000 nM respectively. Immunogenicity of all the screened CTL epitopes was also obtained by using “MHC I Immunogenicity” tool of IEDB (http://tools.iedb.org/immunogenicity/) with all the parameters set to default analyzing 1st, 2nd, and C-terminus amino acids of the given screened epitope [68]. The tool predicts the immunogenicity of a given peptide-MHC (pMHC) complex on the basis of the physiochemical properties of constituting amino acid and their position within the peptide sequence.

##### Screening of Helper T lymphocyte (HTL) Epitopes

To identify the Helper T lymphocyte epitopes from NiV proteins, the IEDB tool “MHC-II Binding Predictions” (http://tools.iedb.org/mhcii/) was used. Peptides with IC50 values <50 nM are considered high affinity, <500 nM intermediate affinity and <5000 nM low affinity [69–72]. The tool generates percentile rank for each potential peptide. This percentile rank is generated by the combination of three different methods viz. combinatorial library, SMM_align and Sturniolo and by comparing the score of the peptide against the scores of other random five million 15-mer peptides of SWISSPROT database [69–72]. The rank from the consensus of all three methods was generated by the median percentile rank of the three methods. The lower the value of percentile the higher would be the rank.

##### Population Coverage by CTL and HTL epitopes

The “Population Coverage” tool of IEDB (http://tools.iedb.org/population/) was used to elucidate the world human population coverage by the shortlisted 33 CTL and 38 HTL epitopes derived from nine NiV proteins [73]. The T cells recognize complex between a specific major MHC molecule and a particular pathogen-derived epitope. The given epitope will elicit a response only in an individual that express an MHC molecule, which is capable of binding that particular epitope. This denominated MHC restriction of T cell responses and the MHC polymorphism provide basis for population coverage study. The MHC types are expressed at dramatically different frequencies in different ethnicities. Hence, a vaccine with optimized population coverage could be of greater relevance in fighting NiV disease [73]. Clinical administration of multiple-epitopes involving both the CTL and the HTL epitopes are predicted here to have a greater probability of larger human population coverage worldwide.

#### B Cell Epitope Prediction

##### Sequence-based B Cell epitope prediction

Protein sequence-based six different methods were utilized to screen linear B cell epitopes from nine different NiV proteins. These methods are available at “B Cell Epitope Prediction Tools” of IEDB server (http://tools.iedb.org/bcell/). In this screening the parameters such as hydrophilicity, flexibility, accessibility, turns, exposed surface, polarity and antigenic propensity of the polypeptides are correlated with their localization in the protein. This allows the search for continuous epitopes prediction from protein sequence. The prediction is based on the propensity scales for each of the 20 amino acids. For a window size n, the i - (n-1)/2 neighboring residues on each side of residue i are used to compute the score for the residue i [74–79]. These methods utilize the propensity scale method as well as the physiochemical properties of the given antigenic sequence to screen potential epitopes using “Bepipred Linear Epitope Prediction”, “Chou & Fasman Beta-Turn Prediction”, “Emini Surface Accessibility Prediction”, “Karplus & Schulz Flexibility Prediction”, “Kolaskar & Tongaonkar Antigenicity” and “Parker Hydrophilicity Prediction” tools [74–79].

##### Structure-based B cell epitope prediction

The Ellipro (ElliPro: Antibody Epitope Prediction tool; http://tools.iedb.org/ellipro/) and the DiscoTope2.0 (DiscoTope: Structure-based Antibody Prediction tool; http://tools.iedb.org/discotope/) methods available at IEDB, were used to screen the linear and the discontinuous B cell epitopes [80, 81]. The ElliPro method analyses on the basis of the location of residue in the protein’s 3D structure. The residues lying outside of an ellipsoid covering 90% of the inner core protein residues score highest Protrusion Index (PI) of 0.9; and so on. The discontinuous epitopes predicted by the ElliPro tool are clustered on the basis of the distance “R” in Å between two residue’s centers of mass lying outside of the largest possible ellipsoid. The larger value of R indicates larger distant residues (residue discontinuity) are screened in the epitopes. The Discotope 2.0 method is based on the “contact number” of the residue’s Cα carbon atom as well as on the propensity of a given residue to be a part of an epitope [80, 81]. The residue “contact number” is the number of Cα atoms in the antigen within a distance of 10 Å of the particular residue’s Cα atom. A low contact number would indicate the residue being close to the surface or in protruding regions of the antigen’s structures.

### Characterisation of potential epitopes

#### Epitope conservation analysis

The shortlisted CTL, HTL and B cell epitopes screened from nine NiV proteins were analysed for the conservancy of their amino acid sequence by “Epitope Conservancy Analysis” tool (http://tools.iedb.org/conservancy/) of IEDB. The epitope conservancy is the fraction of protein sequences that contain that particular epitope. The analysis was done against their entire respective source protein sequences of NiV proteins retrieved from the NCBI protein database [82].

#### Epitope Toxicity prediction

The tool ToxinPred (http://crdd.osdd.net/raghava/toxinpred/multi_submit.php) was used to analyse the toxicity of shortlisted CTL, HTL and B cell epitopes. The tool allows to identifying highly toxic or non-toxic short peptides. The toxicity check analysis was done by the “SVM (Swiss-Prot) based” (support vector machine) method utilizing dataset of 1805 sequences as positive, 3593 negative sequences from Swissprot as well as an alternative dataset comprises the same 1805 positive sequences and 12541 negative sequences from TrEMBLE [83].

#### Overlapping residue analysis

The overlapping residue analysis for the shortlisted CTL, HTL and the B cell linear epitopes was performed by the Multiple Sequence Alignment (MSA) analysis by Clustal Omega tool (https://www.ebi.ac.uk/Tools/msa/clustalo/) of EBI (European Bioinformatics Institute) [84]. The Clustal Omega multiple sequence alignment tool virtually aligns any number of protein sequences and delivers an accurate alignment.

#### Epitope selected for molecular interaction study with HLA allele and TAP transporter

On the basis of the overlapping residue analysis of shortlisted CTL, HTL and linear B cell epitopes few numbers of CTL and HTL epitopes were chosen for further analysis involving stable interaction with their respective HLA allele binders (homology models) and TAP cavity interaction (Supplementary table S2, S3 & S4, Figure 2). These epitopes were chosen on the basis of them having partial or full overlapping sequence region amongst all three types of epitopes (CTL, HTL and B Cell), or having full sequence overlap amongst any of the two types of epitopes, or having the highest number of the HLA allele binders.

### Molecular interaction analysis of selected epitopes with HLA allele and TAP transporter

#### Tertiary structure modelling of HLA alleles and selected T cell epitopes

The Swiss-model was used for homology modelling of the HLA class I and II allele binders of shortlisted epitopes [64]. The amino acid sequence of the HLA allele binders were retrieved from Immuno Polymorphism Database (IPD-IMGT/HLA) (https://www.ebi.ac.uk/ipd/imgt/hla/allele.html). Templates for homology modelling were chosen on the basis of highest amino acid sequence similarity. All the HLA allele models were further validated by their QMEAN value. The QMEAN value gives a composite quality estimate involving both global as well as local analysis of the model [85]. Generated models having acceptable QMEAN value (cutoff −4.0) were chosen for further studies (Supplementary table S2).

The “Natural Peptides Module for Beginners” module of PEPstrMOD (http://osddlinux.osdd.net/raghava/pepstrmod/nat_ss.php) was utilized to generate tertiary structures for the selected CTL and HTL epitopes [86]. The time window for simulation was set to 100 picoseconds (ps) in a vacuum environment.

#### Molecular interaction analysis of selected CTL and HTL epitopes with HLA alleles

The AutoDock 4.2 (ADT) and the AutoDock Vina were used for *in-silico* molecular docking study of the selected CTL and HTL epitopes with their respective HLA class I and II allele binders [87, 88]. The generated docked complexes were studied for their stable nature by molecular dynamics simulation. MD simulation was performed by the Gromacs 5.1.4 using the Optimized Potential for Liquid Simulations - all-atom force field (OPLS-AA) [89, 90].

#### Molecular interaction analysis of selected CTL epitopes with TAP transporter

TAP transporter plays an important role in the presentation of CTL epitopes. Following the immuno-proteasomal processing of xenobiotic proteins, the fragmented peptides get transported to the endoplasmic reticulum (ER) through the TAP transporter. Finally these short peptides reach to the Golgi apparatus from which they get presented on the cell surface [91]. Molecular interaction study of the shortlisted CTL epitopes with the TAP transporter cavity was performed by molecular docking using AutoDock Vina [87, 88]. As structural model the cryo-EM structure of TAP transporter (PDB ID: 5u1d) after removing the antigen from TAP cavity of the original structure [56] was used for epitope-TAP interaction study.

#### Characterisation of designed Multi-Epitope Vaccines

##### Interferon gamma inducing epitope prediction

From the designed amino acid sequence of both the MEVs potential interferon gamma (IFN-γ) epitopes were screened via the “IFNepitope” server (http://crdd.osdd.net/raghava/ifnepitope/scan.php) using the “Motif and SVM hybrid”, (MERCI: Motif-EmeRging and with Classes-Identification, and SVM: support vector machine) method. The tool predicts peptides from protein sequences having the capacity to induce IFN-gamma release from CD4+T cells. This module generates overlapping IFN-gamma inducing peptides from query sequence. For the screening, IEDB database with 3705 IFN-gamma inducing and 6728 non-inducing MHC class II binders is utilized [92, 93].

##### MEVs allergenicity and antigenicity prediction

The designed MEVs were further analysed for allergenicity and antigenicity prediction by utilizing the AlgPred (http://crdd.osdd.net/raghava/algpred/submission.html) and the Vaxigen (http://www.ddg-pharmfac.net/vaxijen/VaxiJen/VaxiJen.html) tools respectively [94, 95]. The AlgPred prediction compares the sequence similarity between known epitopes with any region of the submitted protein. For the screening of allergenicity, the Swiss-prot dataset consisting of 101725 non-allergens and 323 allergens is utilized [94]. The VaxiJen performs an alignment-free approach, solely based on comparing the physicochemical properties of the query amino acid sequence with the Swiss-prot dataset. For prediction of antigenicity, the Bacterial, viral and the tumour protein datasets were used to derive models for the prediction of whole protein antigenicity. Every set consisted of known 100 antigens and 100 non-antigens [95].

##### Physicochemical property analysis of designed MEVs

The ProtParam (https://web.expasy.org/protparam/) tool was utilized to analyse the physiochemical properties of the designed CTL and HTL MEVs [96]. The ProtParam analysis performs an empirical investigation for the given query amino acid sequence. ProtParam computes various physicochemical properties derived from a given protein sequence.

##### Tertiary structure modelling and refinement of MEVs

The tertiary structure of the designed CTL and HTL MEVs were calculated by homology modelling utilizing the RaptorX structure prediction tool (http://raptorx.uchicago.edu/StructurePrediction/predict/) [97]. RaptorX predicts template-based secondary and tertiary structures, protein-protein contacts, solvent accessibility, disordered regions and binding sites for given protein sequence, having close or even less similar homologs in the Protein Data Bank (PDB). RaptorX also assigns confidence scores to indicate the quality of a predicted 3D model [98–100]. Quality assessment for both the generated homology models of CTL and HTL MEVs was performed by their respective P-values. The P-value for a predicted homology model is a probability score of the generated model being worse than the best. Hence the P-value indicates a relative quality of the generated model in terms of modelling error, combining the global distance test (GDT) and the un-normalized global distance test (uGDT) indicating the error involved at each residue. The smaller the P-value the greater the quality of a predicted model.

The refinement of both the generated MEV models was performed by ModRefiner (https://zhanglab.ccmb.med.umich.edu/ModRefiner/) and GalaxyRefine tool (http://galaxy.seoklab.org/cgi-bin/submit.cgi?type=REFINE) [101, 102]. Modrefiner improves the physical realism and structural accuracy of the model using a two-step atomic-level energy minimization. ModRefiner is an algorithm for the atomic-level, high-resolution protein tertiary structure refinement. Both the side-chain and the backbone atoms of a used protein structure are flexible during the structure refinement simulations. The conformational search is guided by a composite of physics and knowledge based force field. The tool generates significant improvement in the physical quality of the local structures [101]. TM-score generated by ModRefiner indicates the structural similarity of the refined model with the original input model. Closer the TM-Score to 1, higher would be the similarity of original and the refined model. The GalaxyRefine tool refines the query tertiary structure by repeated structure perturbation as well as by utilizing the subsequent structural relaxation by the molecular dynamics simulation. The tool GalaxyRefine generates reliable core structures from multiple templates and then re-builds unreliable loops or termini by using an optimization-based refinement method [100, 103]. To avoid any breaks in the 3D model GalaxyRefine uses triaxial loop closure method. The MolProbity score generated for a given refined model indicates the log-weighted combination of the clash score (the number of atomic clashes per 1000 atoms), the Ramachandran favored backbone torsion angles and the percentage of bad side-chain rotamers (the percentages of rotamer outliers). The ‘GDT-HA’ (Global Distance Test-High Accuracy) generated by the tool indicates the backbone structure accuracy; ‘RMSD’ (Root mean Square Deviation) indicates the overall structural deviation in refined model from the initial model and the ‘Rama favored’ indicates percentage of Ramachandran favored residues.

##### Validation of CTL and HTL MEVs refined models

Both the refined CTL and HTL MEV 3D models were further validated by RAMPAGE analysis tool (http://mordred.bioc.cam.ac.uk/~rapper/rampage.php) [104, 105]. The generated Ramachandran plots for the MEV models show the sterically allowed and disallowed residues along with their dihedral psi (ψ) and phi (φ) angles.

##### Discontinuous B-cell epitope prediction from MEVs

Both the generated tertiary models of designed CTL and HTL MEVs were subjected to discontinuous B cell epitopes prediction. The structure-based discontinuous B cell epitopes were screened from both the MEV models by utilizing the ElliPro method as described earlier [80].

#### Molecular interaction analysis of MEVs with TLR-3

Molecular interaction analysis of both the designed MEVs with TLR-3, was performed by molecular docking and molecular dynamics simulation. Molecular docking was performed by PatchDock server (http://bioinfo3d.cs.tau.ac.il/PatchDock/) [106–108]. PatchDock utilizes algorithm for unbound (real life) docking of molecules for protein-protein. The algorithm carries out the rigid docking, with the surface variability/flexibility implicitly addressed through liberal intermolecular penetration. The algorithm focuses on the (i) initial molecular surface fitting on localized, curvature based surface patches (ii) use of Geometric Hashing and Pose Clustering for initial transformation detection (iii) computation of shape complementarity utilizing the Distance Transform (iv) efficient steric clash detection and geometric fit scoring based on a multi-resolution shape representation and (v) utilization of biological information by focusing on hot spot rich surface patches [106–108]. For molecular docking, the 3D structure of the human TLR-3 ectodomain (ECD) was retrieved from the PDB databank (PDB ID: 2A0Z). Further, the molecular dynamics simulation study of the MEVs-TLR-3 complexes was performed by YASARA tool (Yet Another Scientific Artificial Reality Application) [109]. The MD simulations studies were carried out in an explicit water environment in a dodecahedron simulation box at a stabilized temperature of 298K, the pressure of 1atm and pH 7.4, with periodic cell boundary condition. The solvated systems were neutralized with counter ions (NaCl) (concentration 0.9 M). The AMBER14 force field was used on the systems during MD simulation [110]. The Long-range electrostatic energy and forces were calculated using particle-mesh-based Ewald method [111]. The solvated structures were energy minimized by the steepest descent method at a temperature of 298K and a stable pressure of 1 atm. Further, the complexes were equilibrated for period of 1 ns. After equilibration, a production MD simulation was run for 100 ns at a stable temperature and pressure and time frames were saved at every 100 ps, for each MD simulations. The RMSD and RMSF values for Cα, Backbone and all the atoms of all the CTL and HTL MPVs in complex with TLR3 were analyzed for each MD simulation study.

### *In-silico* analysis of MEVs for cloning and expression

*Analysis of cDNA of the MEVs for cloning and expression in mammalian cell line.* Complementary DNA of both the MEVs, codon optimized for expression in mammalian cells were generated by Java Codon Adaptation Tool (http://www.jcat.de/). The generated cDNA was further analysed with the GenScript Rare Codon Analysis Tool (https://www.genscript.com/tools/rare-codon-analysis) which calculates the GC content, Codon Adaptation Index (CAI) and the Tandem rare codon frequency for a given cDNA [112–114]. The CAI indicates the possibility of cDNA expression in a chosen expression system. The tandem rare codon frequency indicates the presence of low-frequency codons in the given cDNA.

## ACKNOWLEDGEMENTS

We acknowledge the Indian Foundation for Fundamental Research Trust (IFFR Trust) for providing resources and funding

## AUTHOR CONTRIBUTION

Idea conceived, methodology designed and performed by S.S., MD simulation done by S.V., critical data analysis and scientific writing was done by S.S., M.K. and M.Kolbe, draft was finalized by S.S., S.V., M.K., M.Kolbe, A.K.S. and K.C.P.

## ADDITIONAL INFORMATION

Authors declare to have no competing interests.

**Supplementary figure S1.**
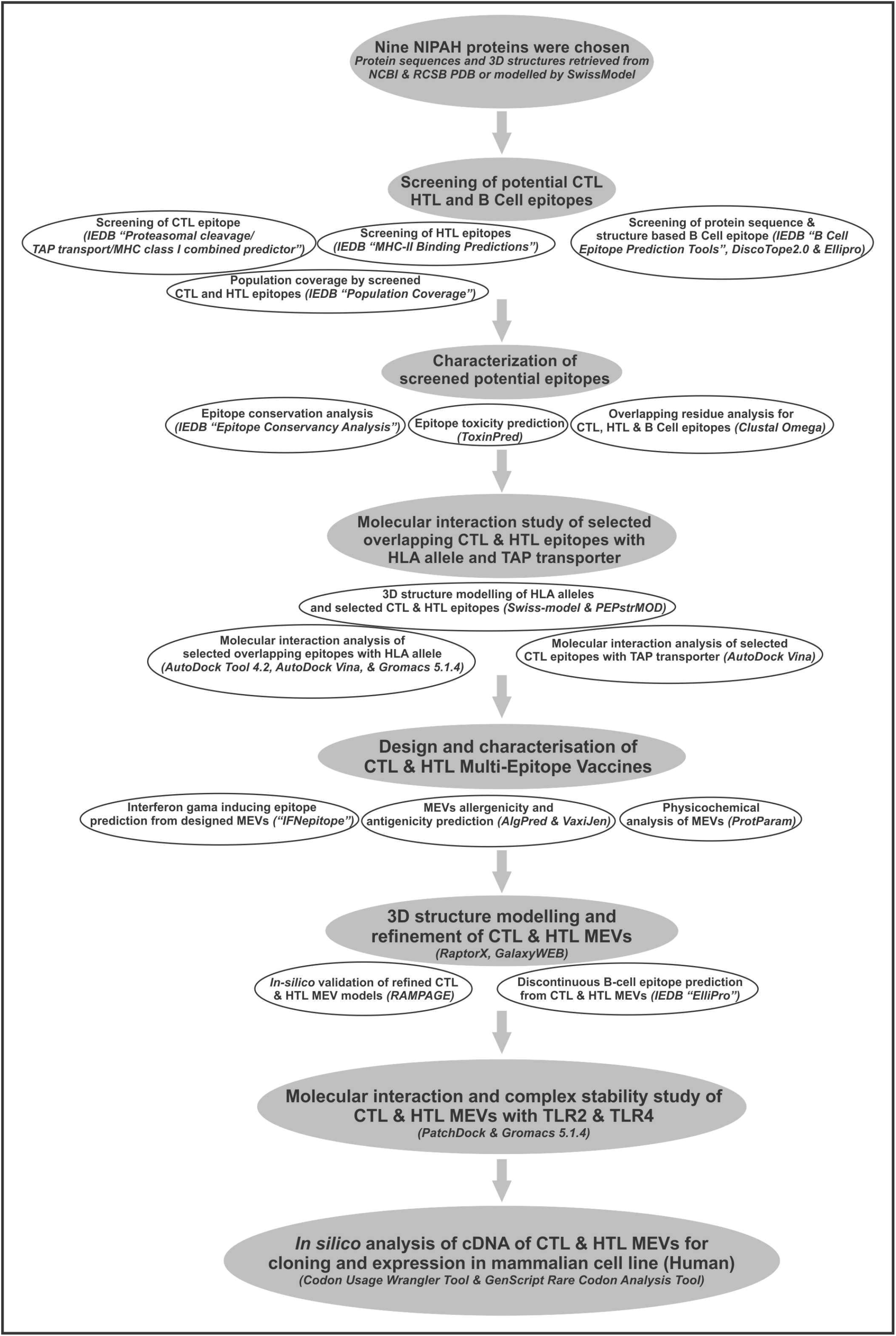
Workflow chart.

**Supplementary figure S2.**
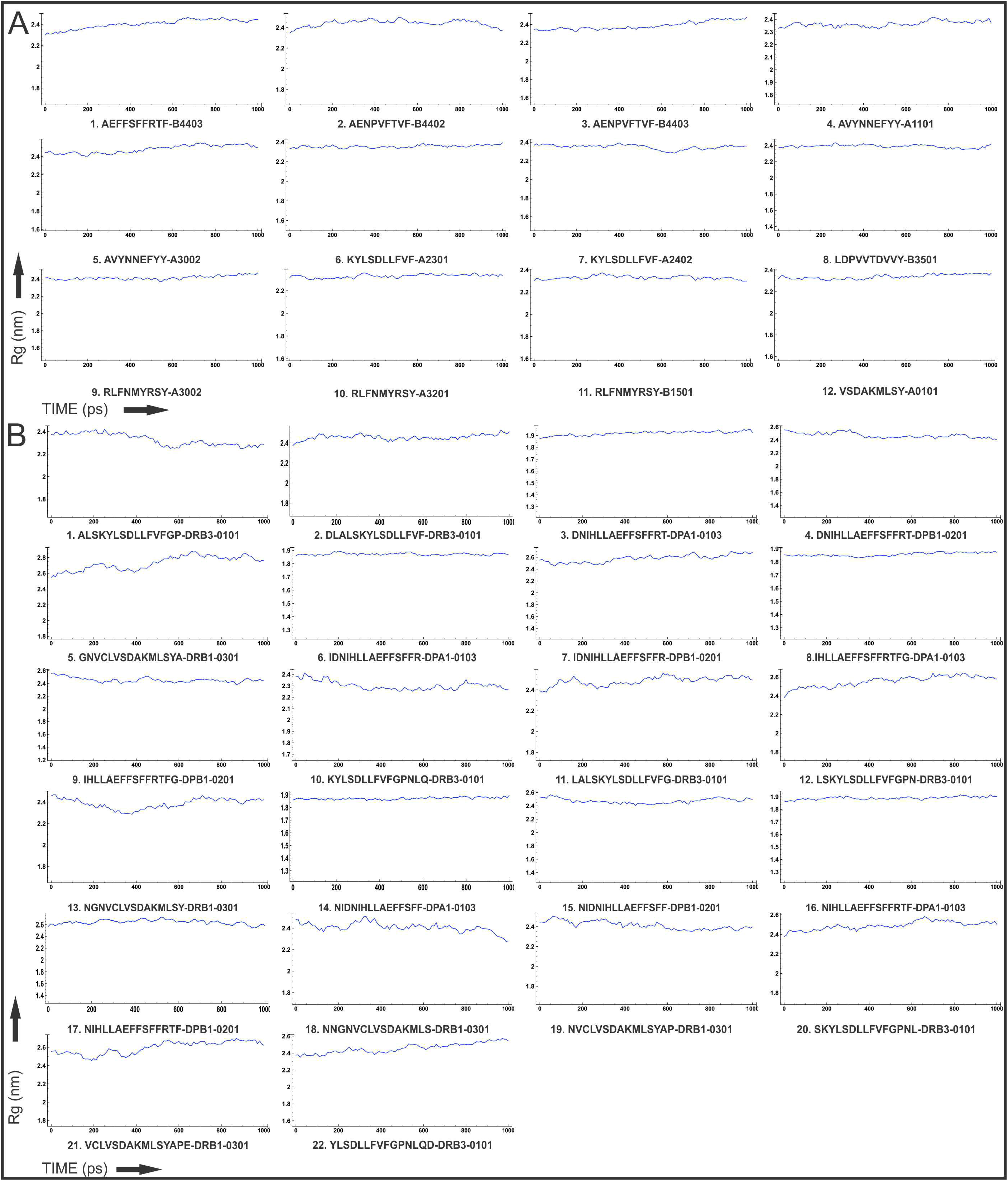
**(A)** Rg (radius of gyration) for the CTL epitope – HLA class I allele complexes, across the time window of 1 nano second. **(B)** Rg for the HTL epitope – HLA class II allele complexes, across the time window of 1 nano second.

**Supplementary figure S3.**
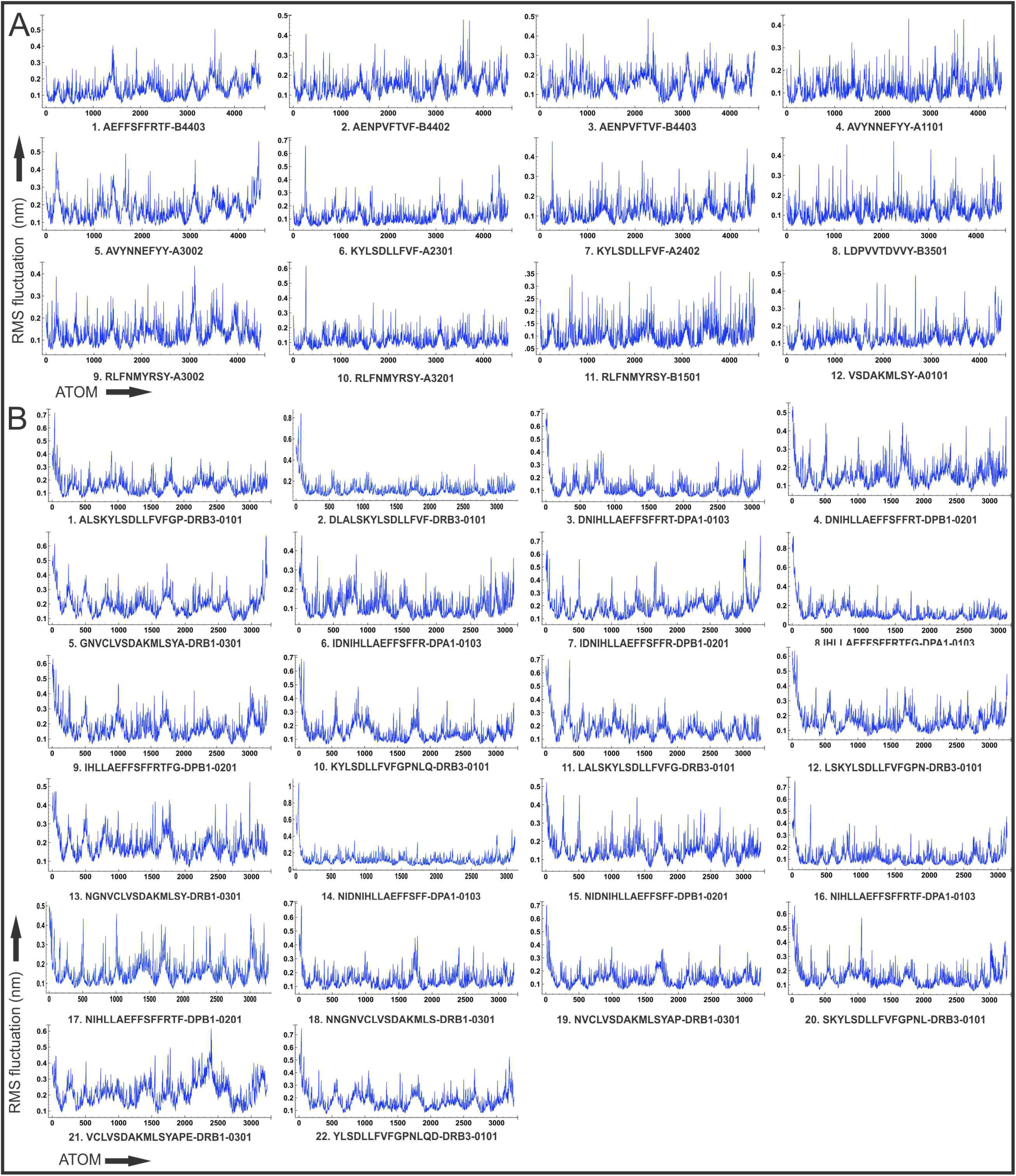
**(A)** RMS fluctuation in nanometers for all the atoms of the CTL epitope – HLA class I allele complexes. **(B)** RMS fluctuation in nanometers for all the atoms of the HTL epitope – HLA class II allele complexes.

**Supplementary figure S4.**
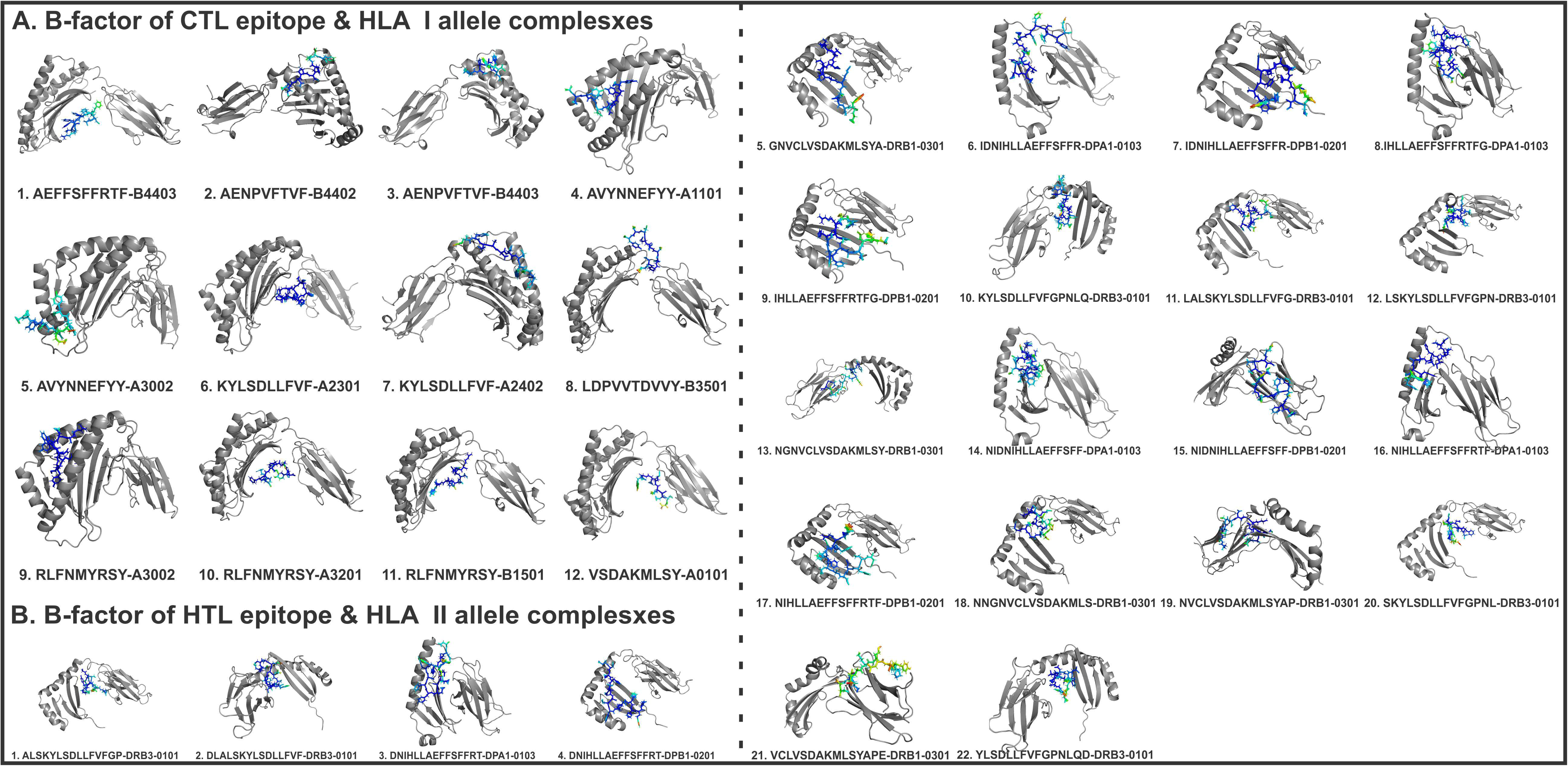
**(A)** B-Factor of CTL epitope – HLA class I allele complexes **(B)** B-Factor of HTL epitope – HLA class II allele complexes. Epitopes are shown in sticks and HLA alleles are shown in gray cartoons. B-factor is indicated by rainbow (VIBGYOR) colour, blue for stable region and red for most unstable region.

**Supplementary table S1. Protein sequence retrieval, tertiary structures retrieval and homology modeling of nine Nipah proteins.** Nipah protein sequences were retrieved from NCBI. Available structure files (pdb) for Nipah proteins were retrieved from RCSB PDB. Nipah proteins with no tertiary structure available were subjected to homology modeling by Swissmodel.

**Supplementary table S2. Homology modeling for HLA alleles.** Tertiary structure of HLA alleles were modeled by homology modeling using SwissModel server. Templates were chosen with highest sequence identity. Generated models with acceptable QMEAN value were chosen for further studies.

**Supplementary table S3. Shortlisted high scoring CTL epitopes.** Selected high scoring CTL epitopes and their respective HLA alleles binders are listed. *In-silico* analysis have shown all the selected epitopes to be non-toxic (Non-Toxin) as well as they show significant conservancy. ToxinPred analysis is based on the ToxinPred main dataset used by “ToxinPred” algorithm to predict toxicity of any unknown peptide. # Epitope match with previous studies indicating consensus in epitope screening by different approaches and methods.

**Supplementary table S4. Shortlisted high scoring HTL epitopes.** Selected high scoring HTL epitopes and their respective HLA alleles binders are listed above. *In-silico* analysis have shown all the selected epitopes to be non-toxic (Non-Toxin) as well as they show significant conservancy.

**Supplementary table S5. World population coverage by the shortlisted CTL and HTL epitopes combined.** The cumulative percent of world (countries as listed in table) population coverage is 97.88% by the joint administration of selected CTL and HTL epitopes as vaccine candidate.

**Supplementary table S6. Shortlisted B Cell epitopes.** BepiPred Linear B Cell epitopes showing sequence overlap with CTL and HTL epitopes are shortlisted above. *In-silico* analysis have shown all the selected epitopes to be non-toxic (Non-Toxin) as well as they show significant amino acid sequence conservancy. #Epitope match with previous studies indicating consensus in epitope screening by different approaches and methods.

**Supplementary table S7. CTL epitope prediction.** Detailed scoring of all screened CTL epitopes and their respective HLA class I allele binders. CTL epitopes were chosen on the basis of high “Total score” and higher number of HLA allele binders. Total score is a combined score of TAP score, MHC score, Proteasome score and Processing score.

**Supplementary table S8. HTL epitope prediction.** Percentile rank of HTL epitopes and their respective HLA class II allele binders. HTL epitopes were screened on the basis of percentile rank (lower the percentile number, higher the rank) and larger number of HLA allele binders. Last column show the method used for epitope screening.

**Supplementary table S9. INF-**γ **epitopes from CTL and HTL MEVs.** INF-γ inducing (POSITIVE) epitopes from CTL and HTL MEVs were screened by using “Motif and SVM hybrid” (MERCI & SVM) approaches.

**Supplementary table S10. Refinement models of CTL and HTL MEVs.** CTL and HTL MEVs models were refined by GalaxyWEB server and used for further studies. After refinement in particular Rama favored residues increased significantly.

**Supplementary table S11. B Cell discontineous epitopes of CTL & HTL MEVs.** Discontinuous B Cell epitopes predicted by ElliPro (IEDB) from CTL & HTL MEVs.

## REFERENCES

1. Angeletti, S., Presti, A.L., Cella, E. and Ciccozzi, M., 2016. Molecular epidemiology and phylogeny of nipah virus infection: a mini review. Asian Pacific journal of tropical medicine, 9(7), pp.630–634.

2. Aguilar, H.C., Henderson, B.A., Zamora, J.L. and Johnston, G.P., 2016. Paramyxovirus glycoproteins and the membrane fusion process. Current clinical microbiology reports, 3(3), pp.142–154.

3. Ang, B.S., Lim, T.C. and Wang, L., 2018. Nipah Virus Infection. Journal of clinical microbiology, pp.JCM-01875.

4. WHO Report, Surveillance and outbreak alert, Nipah virus; https://www.who.int/health-topics/nipah-virus-infection#tab=tab_1

5. Plowright RK, Becker DJ, Crowley DE, Washburne AD, Huang T, Nameer PO, Gurley ES, Han BA. Prioritizing surveillance of Nipah virus in India. PLoS Negl Trop Dis. 2019 Jun 27;13(6):e0007393. doi: 10.1371/journal.pntd.0007393. PMID: 31246966; PMCID: PMC6597033.

6. Thomas, B., Chandran, P., Lilabi, M. P., George, B., Sivakumar, C. P., Jayadev, V. K., Bindu, V., Rajasi, R. S., Vijayan, B., Mohandas, A., & Hafeez, N. (2019). Nipah Virus Infection in Kozhikode, Kerala, South India, in 2018: Epidemiology of an Outbreak of an Emerging Disease. Indian journal of community medicine : official publication of Indian Association of Preventive & Social Medicine, 44(4), 383–387. https://doi.org/10.4103/ijcm.IJCM_198_19

7. Mathieu, C., Guillaume, V., Volchkova, V.A., Pohl, C., Jacquot, F., Looi, R.Y., Wong, K.T., Legras-Lachuer, C., Volchkov, V.E., Lachuer, J. and Horvat, B., 2012. Nonstructural Nipah virus C protein regulates both the early host proinflammatory response and viral virulence. Journal of virology, pp.JVI-01203.

8. Liu, Q., Chen, L., Aguilar, H.C. and Chou, K.C., 2018. A stochastic assembly model for Nipah virus revealed by super-resolution microscopy. Nature communications, 9(1), p.3050.

9. Johnston, G.P., Contreras, E.M., Dabundo, J., Henderson, B.A., Matz, K.M., Ortega, V., Ramirez, A., Park, A. and Aguilar, H.C., 2017. Cytoplasmic motifs in the nipah virus fusion protein modulate virus particle assembly and egress. Journal of virology, pp.JVI-02150.

10. Satterfield, B.A., Cross, R.W., Fenton, K.A., Borisevich, V., Agans, K.N., Deer, D.J., Graber, J., Basler, C.F., Geisbert, T.W. and Mire, C.E., 2016. The Nipah virus C and W proteins contribute to respiratory disease in ferrets. Journal of virology, pp.JVI-00215.

11. Lamp, B., Dietzel, E., Kolesnikova, L., Sauerhering, L., Erbar, S., Weingartl, H. and Maisner, A., 2013. Nipah virus entry and egress from polarized epithelial cells. Journal of virology, pp.JVI-02696.

12. Weise, C., Erbar, S., Lamp, B., Vogt, C., Diederich, S. and Maisner, A., 2010. Tyrosine residues in the cytoplasmic domains affect sorting and fusion activity of the Nipah virus glycoproteins in polarized epithelial cells. Journal of virology, 84(15), pp.7634–7641.

13. Ciancanelli, M.J. and Basler, C.F., 2006. Mutation of YMYL in the Nipah virus matrix protein abrogates budding and alters subcellular localization. Journal of virology, 80(24), pp.12070–12078.

14. Patch, J.R., Crameri, G., Wang, L.F., Eaton, B.T. and Broder, C.C., 2007. Quantitative analysis of Nipah virus proteins released as virus-like particles reveals central role for the matrix protein. Virology journal, 4(1), p.1.

15. Patch, J.R., Han, Z., McCarthy, S.E., Yan, L., Wang, L.F., Harty, R.N. and Broder, C.C., 2008. The YPLGVG sequence of the Nipah virus matrix protein is required for budding. Virology journal, 5(1), p.137.

16. Jordan, P.C., Liu, C., Raynaud, P., Lo, M.K., Spiropoulou, C.F., Symons, J.A., Beigelman, L. and Deval, J., 2018. Initiation, extension, and termination of RNA synthesis by a paramyxovirus polymerase. PLoS pathogens, 14(2), p.e1006889.

17. Ranadheera, C., Proulx, R., Chaiyakul, M., Jones, S., Grolla, A., Leung, A., Rutherford, J., Kobasa, D., Carpenter, M. and Czub, M., 2018. The interaction between the Nipah virus nucleocapsid protein and phosphoprotein regulates virus replication. Scientific reports, 8(1), p.15994.

18. Baronti, L., Erales, J., Habchi, J., Felli, I.C., Pierattelli, R. and Longhi, S., 2015. Dynamics of the intrinsically disordered C-terminal domain of the Nipah virus nucleoprotein and interaction with the X domain of the phosphoprotein as unveiled by NMR spectroscopy. ChemBioChem, 16(2), pp.268–276.

19. Uchida, S., Horie, R., Sato, H., Kai, C. and Yoneda, M., 2018. Possible role of the Nipah virus V protein in the regulation of the interferon beta induction by interacting with UBX domain-containing protein1. Scientific reports, 8(1), p.7682.

20. Ludlow, L.E., Lo, M.K., Rodriguez, J.J., Rota, P.A. and Horvath, C.M., 2008. Henipavirus V protein association with Polo-like kinase reveals functional overlap with STAT1 binding and interferon evasion. Journal of virology, 82(13), pp.6259–6271.

21. Park, M.S., Shaw, M.L., Munoz-Jordan, J., Cros, J.F., Nakaya, T., Bouvier, N., Palese, P., García-Sastre, A. and Basler, C.F., 2003. Newcastle disease virus (NDV)-based assay demonstrates interferon-antagonist activity for the NDV V protein and the Nipah virus V, W, and C proteins. Journal of virology, 77(2), pp.1501–1511.

22. Sakib, M.S., Islam, M., Hasan, A.K.M. and Nabi, A.H.M., 2014. Prediction of epitope-based peptides for the utility of vaccine development from fusion and glycoprotein of nipah virus using in silico approach. Advances in bioinformatics, 2014.

23. Guillaume, V., Contamin, H., Loth, P., Georges-Courbot, M.C., Lefeuvre, A., Marianneau, P., Chua, K.B., Lam, S.K., Buckland, R., Deubel, V. and Wild, T.F., 2004. Nipah virus: vaccination and passive protection studies in a hamster model. Journal of virology, 78(2), pp.834–840.

24. Koyuncu, O.O., Hogue, I.B. and Enquist, L.W., 2013. Virus infections in the nervous system. Cell host & microbe, 13(4), pp.379–393.

25. Griffin, D.E. and Metcalf, T., 2011. Clearance of virus infection from the CNS. Current opinion in virology, 1(3), pp.216–221.

26. Kong, D., Wen, Z., Su, H., Ge, J., Chen, W., Wang, X., Wu, C., Yang, C., Chen, H. and Bu, Z., 2012. Newcastle disease virus-vectored Nipah encephalitis vaccines induce B and T cell responses in mice and long-lasting neutralizing antibodies in pigs. Virology, 432(2), pp.327–335.

27. Kamthania, M. and Sharma, D.K., 2015. Screening and structure-based modeling of T-cell epitopes of Nipah virus proteome: an immunoinformatic approach for designing peptide-based vaccine. 3 Biotech, 5(6), pp.877-882.

28. Kamthania, M. and Sharma, D.K., 2016. Epitope-based peptides prediction from proteome of nipah virus. International Journal of Peptide Research and Therapeutics, 22(4), pp.465–470.

29. Ali, M.T., Morshed, M.M. and Hassan, F., 2015. A computational approach for designing a universal epitope-based peptide vaccine against Nipah virus. Interdisciplinary Sciences: Computational Life Sciences, 7(2), pp.177–185.

30. kumar Sharma, S., Srivastava, S., Kumar, A. and Srivastava, V., 2021. Anticipation of Antigenic Sites for the Goal of Vaccine Designing Against Nipah Virus: An Immunoinformatics Inquisitive Quest. International Journal of Peptide Research and Therapeutics, pp.1–13.

31. Dey, S., Roy, P., Dutta, T., Nandy, A. and Basak, S.C., 2018. Rational Design of Peptide Vaccines for the Highly Lethal Nipah and Hendra Viruses. bioRxiv, p.425819.

32. Krishnamoorthy, P.K., Subasree, S., Arthi, U., Mobashir, M., Gowda, C. and Revanasiddappa, P.D., 2020. T-cell Epitope-based Vaccine Design for Nipah Virus by Reverse Vaccinology Approach. Combinatorial chemistry & high throughput screening, 23(8), pp.788–796.

33. Sakib, M.S., Islam, M., Hasan, A.K.M. and Nabi, A.H.M., 2014. Prediction of epitope-based peptides for the utility of vaccine development from fusion and glycoprotein of nipah virus using in silico approach. Advances in bioinformatics, 2014.

34. Eshaghi, M., Tan, W.S. and Yusoff, K., 2005. Identification of epitopes in the nucleocapsid protein of Nipah virus using a linear phage-displayed random peptide library. Journal of medical virology, 75(1), pp.147–152.

35. Mohammed, A.A., Shantier, S.W., Mustafa, M.I., Osman, H.K., Elmansi, H.E., Osman, I.A.A., Mohammed, R.A., Abdelrhman, F.A., Elnnewery, M.E., Yousif, E.M. and Mustafa, M.M., 2020. Epitope-based peptide vaccine against glycoprotein G of Nipah henipavirus using immunoinformatics approaches. Journal of immunology research, 2020.

36. Gupta, A.K., Kumar, A., Rajput, A., Kaur, K., Dar, S.A., Thakur, A., Megha, K. and Kumar, M., 2020. NipahVR: a resource of multi-targeted putative therapeutics and epitopes for the Nipah virus. Database, 2020.

37. Habib, P.T., 2021. Learning from COVID-19 Pandemic: In Silico Vaccine and Cloning Design Against Nipah Virus by Studying and Analyzing the Whole Nipah Virus Proteome.

38. Singh, R.K., Dhama, K., Chakraborty, S., Tiwari, R., Natesan, S., Khandia, R., Munjal, A., Vora, K.S., Latheef, S.K., Karthik, K. and Singh Malik, Y., 2019. Nipah virus: epidemiology, pathology, immunobiology and advances in diagnosis, vaccine designing and control strategies–a comprehensive review. Veterinary Quarterly, 39(1), pp.26–55.

39. Majee, P., Jain, N. and Kumar, A., 2021. Designing of a multi-epitope vaccine candidate against Nipah virus by in silico approach: a putative prophylactic solution for the deadly virus. Journal of Biomolecular Structure and Dynamics, 39(4), pp.1461–1480.

40. Ojha, R., Pareek, A., Pandey, R.K., Prusty, D. and Prajapati, V.K., 2019. Strategic development of a next-generation multi-epitope vaccine to prevent Nipah virus zoonotic infection. ACS omega, 4(8), pp.13069–13079.

41. Majee, P., Jain, N. and Kumar, A., 2021. Designing of a multi-epitope vaccine candidate against Nipah virus by in silico approach: a putative prophylactic solution for the deadly virus. Journal of Biomolecular Structure and Dynamics, 39(4), pp.1461–1480.

42. Mohammed, A.A., Shantier, S.W., Mustafa, M.I., Osman, H.K., Elmansi, H.E., Osman, I.A.A., Mohammed, R.A., Abdelrhman, F.A., Elnnewery, M.E., Yousif, E.M. and Mustafa, M.M., 2020. Epitope-based peptide vaccine against glycoprotein G of Nipah henipavirus using immunoinformatics approaches. Journal of immunology research, 2020.

43. Ali, M.T., Morshed, M.M. and Hassan, F., 2015. A computational approach for designing a universal epitope-based peptide vaccine against Nipah virus. Interdisciplinary Sciences: Computational Life Sciences, 7(2), pp.177–185.

44. Soltan, M.A., Eldeen, M.A., Elbassiouny, N., Mohamed, I., El-Damasy, D.A., Fayad, E., Abu Ali, O.A., Raafat, N., Eid, R.A. and Al-Karmalawy, A.A., 2021. Proteome Based Approach Defines Candidates for Designing a Multitope Vaccine against the Nipah Virus. International Journal of Molecular Sciences, 22(17), p.9330.

45. Wilson, S.S., Wiens, M.E. and Smith, J.G., 2013. Antiviral mechanisms of human defensins. Journal of molecular biology, 425(24), pp.4965–4980.

46. Duits, L.A., Nibbering, P.H., Strijen, E., Vos, J.B., Mannesse-Lazeroms, S.P., Sterkenburg, M.A. and Hiemstra, P.S., 2003. Rhinovirus increases human β-defensin-2 and-3 mRNA expression in cultured bronchial epithelial cells. Pathogens and Disease, 38(1), pp.59–64.

47. Yang, D., Biragyn, A., Kwak, L.W. and Oppenheim, J.J., 2002. Mammalian defensins in immunity: more than just microbicidal. Trends in immunology, 23(6), pp.291–296.

48. Biragyn, A., Surenhu, M., Yang, D., Ruffini, P.A., Haines, B.A., Klyushnenkova, E., Oppenheim, J.J. and Kwak, L.W., 2001. Mediators of innate immunity that target immature, but not mature, dendritic cells induce antitumor immunity when genetically fused with nonimmunogenic tumor antigens. The Journal of Immunology, 167(11), pp.6644–6653.

49. Duits, L.A., Nibbering, P.H., van Strijen, E., Vos, J.B., Mannesse-Lazeroms, S.P., van Sterkenburg, M.A. and Hiemstra, P.S., 2003. Rhinovirus increases human β-defensin-2 and-3 mRNA expression in cultured bronchial epithelial cells. FEMS Immunology & Medical Microbiology, 38(1), pp.59–64.

50. Kohlgraf, K.G., Pingel, L.C., Dietrich, D.E. and Brogden, K.A., 2010. Defensins as anti-inflammatory compounds and mucosal adjuvants. Future microbiology, 5(1), pp.99–113.

51. Hu W, Li F, Yang X, Li Z, Xia H, Li G, Wang Y, Zhang Z. A flexible peptide linker enhances the immunoreactivity of two copies HBsAg preS1 (21-47) fusion protein. J Biotechnol. 2004;107:83–90.

52. Hajighahramani, N., Nezafat, N., Eslami, M., Negahdaripour, M., Rahmatabadi, S.S. and Ghasemi, Y., 2017. Immunoinformatics analysis and in silico designing of a novel multi-epitope peptide vaccine against Staphylococcus aureus. Infection, Genetics and Evolution, 48, pp.83–94.

53. Chen, X., Zaro, J.L. and Shen, W.C., 2013. Fusion protein linkers: property, design and functionality. Advanced drug delivery reviews, 65(10), pp.1357–1369.

54. Hoover, D.M., Rajashankar, K.R., Blumenthal, R., Puri, A., Oppenheim, J.J., Chertov, O. and Lubkowski, J., 2000. The structure of human β-defensin-2 shows evidence of higher order oligomerization. Journal of Biological Chemistry, 275(42), pp.32911–32918. PDB ID: 1FD3.

55. Antoniou, A.N., Powis, S.J. and Elliott, T., 2003. Assembly and export of MHC class I peptide ligands. Current opinion in immunology, 15(1), pp.75–81.

56. Oldham, M.L., Grigorieff, N. and Chen, J., 2016. Structure of the Transporter associated with antigen processing trapped by herpes simplex virus. eLife, 5, p.e21829.

57. Meena, S.R., Gangwar, S.P. and Saxena, A.K., 2012. Purification, crystallization and preliminary X-ray crystallographic analysis of the ATPase domain of human TAP in nucleotide-free and ADP-, vanadate- and azide-complexed forms. Acta Crystallographica Section F: Structural Biology and Crystallization Communications, 68(6), pp.655–658.

58. Delneste, Y., Beauvillain, C. and Jeannin, P., 2007. Innate immunity: structure and function of TLRs. Medecine sciences: M/S, 23(1), pp.67–73.

59. Totura, A.L., Whitmore, A., Agnihothram, S., Schäfer, A., Katze, M.G., Heise, M.T. and Baric, R.S., 2015. Toll-like receptor 3 signaling via TRIF contributes to a protective innate immune response to severe acute respiratory syndrome coronavirus infection. MBio, 6(3), pp.e00638–15.

60. Shaw, M.L., Cardenas, W.B., Zamarin, D., Palese, P. and Basler, C.F., 2005. Nuclear localization of the Nipah virus W protein allows for inhibition of both virus-and toll-like receptor 3-triggered signaling pathways. Journal of virology, 79(10), pp.6078–6088.

61. Seto, J., Qiao, L., Guenzel, C.A., Xiao, S., Shaw, M.L., Hayot, F. and Sealfon, S.C., 2010. Novel Nipah virus immune-antagonism strategy revealed by experimental and computational study. Journal of virology, 84(21), pp.10965–10973.

62. Farina, C., Krumbholz, M., Giese, T., Hartmann, G., Aloisi, F. and Meinl, E., 2005. Preferential expression and function of Toll-like receptor 3 in human astrocytes. Journal of neuroimmunology, 159(1-2), pp.12–19.

63. Weingartl, H., Czub, S., Copps, J., Berhane, Y., Middleton, D., Marszal, P., Gren, J., Smith, G., Ganske, S., Manning, L. and Czub, M., 2005. Invasion of the central nervous system in a porcine host by Nipah virus. Journal of virology, 79(12), pp.7528–7534.

64. Arnold K, Bordoli L, Kopp J, and Schwede T (2006). The SWISS-MODEL Workspace: A web-based environment for protein structure homology modelling. Bioinformatics.,22,195–201.

65. Tenzer S, Peters B, Bulik S, Schoor O, Lemmel C, Schatz MM, Kloetzel PM, Rammensee HG, Schild H, Holzhutter HG. 2005. Modeling the MHC class I pathway by combining predictions of proteasomal cleavage, TAP transport and MHC class I binding. Cell Mol Life Sci 62:1025–1037.

66. Peters B, Bulik S, Tampe R, Van Endert PM, Holzhutter HG. 2003. Identifying MHC class I epitopes by predicting the TAP transport efficiency of epitope precursors. J Immunol171:1741–1749.

67. Hoof, I., Peters, B., Sidney, J., Pedersen, L.E., Sette, A., Lund, O., Buus, S. and Nielsen, M., 2009. NetMHCpan, a method for MHC class I binding prediction beyond humans. Immunogenetics, 61(1), p.1.

68. Calis JJA, Maybeno M, Greenbaum JA, Weiskopf D, De Silva AD, Sette A, Kesmir C, Peters B. 2013. Properties of MHC class I presented peptides that enhance immunogenicity. PloS Comp. Biol. 8(1):361.

69. Wang, P., Sidney, J., Kim, Y., Sette, A., Lund, O., Nielsen, M. and Peters, B., 2010. Peptide binding predictions for HLA DR, DP and DQ molecules. BMC bioinformatics, 11(1), p.568.

70. Sidney, J., Assarsson, E., Moore, C., Ngo, S., Pinilla, C., Sette, A. and Peters, B., 2008. Quantitative peptide binding motifs for 19 human and mouse MHC class I molecules derived using positional scanning combinatorial peptide libraries. Immunome research, 4(1), p.2.

71. Nielsen, M., Lundegaard, C. and Lund, O., 2007. Prediction of MHC class II binding affinity using SMM-align, a novel stabilization matrix alignment method. BMC bioinformatics, 8(1), p.238.

72. Sturniolo, T., Bono, E., Ding, J., Raddrizzani, L., Tuereci, O., Sahin, U., Braxenthaler, M., Gallazzi, F., Protti, M.P., Sinigaglia, F. and Hammer, J., 1999. Generation of tissue-specific and promiscuous HLA ligand databases using DNA microarrays and virtual HLA class II matrices. Nature biotechnology, 17(6), p.555.

73. Bui H. H, Sidney J, Dinh K, Southwood S, Newman M. J, Sette A. 2006. Predicting population coverage of T-cell epitope-based diagnostics and vaccines. BMC Bioinformatics 17:153.

74. Larsen JE, Lund O, Nielsen M. 2006. Improved method for predicting linear B-cell epitopes. Immunome Res 2:2.

75. Chou PY, Fasman GD. 1978. Prediction of the secondary structure of proteins from their amino acid sequence. Adv Enzymol Relat Areas Mol Biol 47:45–148.

76. Emini EA, Hughes JV, Perlow DS, Boger J. 1985. Induction of hepatitis A virus-neutralizing antibody by a virus-specific synthetic peptide. J Virol 55:836–839.

77. Karplus PA, Schulz GE. 1985. Prediction of chain flexibility in proteins. Naturwissenschaften 72:212–213.

78. Kolaskar AS, Tongaonkar PC. 1990. A semi-empirical method for prediction of antigenic determinants on protein antigens. FEBS Lett276:172–174.

79. Parker JM, Guo D, Hodges RS. 1986. New hydrophilicity scale derived from high-performance liquid chromatography peptide retention data: correlation of predicted surface residues with antigenicity and X-ray-derived accessible sites. Biochemistry 25:5425–5432.

80. J. V. Kringelum, C. Lundegaard, O. Lund, M. Nielsen. 2012. Reliable B cell epitope predictions: impacts of method development and improved benchmarking. PLoS Comput Biol. 8:(12):e1002829.

81. Ponomarenko JV, Bui H, Li W, Fusseder N, Bourne PE, Sette A, Peters B. 2008. ElliPro: a new structure-based tool for the prediction of antibody epitopes. BMC Bioinformatics 9:514.

82. Bui HH, Sidney J, Li W, Fusseder N, Sette A. 2007. Development of an epitope conservancy analysis tool to facilitate the design of epitope-based diagnostics and vaccines. BMC Bioinformatics 8:361.

83. Gupta, S., Kapoor, P., Chaudhary, K., Gautam, A., Kumar, R., Raghava, G.P. and Open Source Drug Discovery Consortium, 2013. In silico approach for predicting toxicity of peptides and proteins. PLoS One, 8(9), p.e73957.

84. Sievers, F., Wilm, A., Dineen, D., Gibson, T.J., Karplus, K., Li, W., Lopez, R., McWilliam, H., Remmert, M., Söding, J. and Thompson, J.D., and Higgins D.G., 2011. Fast, scalable generation of high-quality protein multiple sequence alignments using Clustal Omega. Molecular systems biology, 7(1), p.539.

85. Benkert, P., Tosatto, S.C. and Schomburg, D., 2008. QMEAN: A comprehensive scoring function for model quality assessment. Proteins: Structure, Function, and Bioinformatics, 71(1), pp.261–277.

86. Singh, S., Singh, H., Tuknait, A., Chaudhary, K., Singh, B., Kumaran, S. and Raghava, G.P.S. (2015) PEPstrMOD: structure prediction of peptides containing natural, non-natural and modified residues. Biology Direct 10:73.

87. Morris, G.M., Huey, R., Lindstrom, W., Sanner, M.F., Belew, R.K., Goodsell, D.S. and Olson, A.J., 2009. AutoDock4 and AutoDockTools4: Automated docking with selective receptor flexibility. Journal of computational chemistry, 30(16), pp.2785–2791.

88. O. Trott, A. J. Olson, AutoDock Vina: improving the speed and accuracy of docking with a new scoring function, efficient optimization and multithreading, Journal of Computational Chemistry 31 (2010) 455–461

89. Abraham, M. J., Murtola, T., Schulz, R., Pa ll, S., Smith, J. C., Hess, B., Lindahl, E. GROMACS: High performance molecular simulations through multi-level parallelism from lap-tops to supercomputers. SoftwareX 1– 2:19–25, 2015.

90. Jorgensen, W.L., Maxwell, D.S. and Tirado-Rives, J., 1996. Development and testing of the OPLS all-atom force field on conformational energetics and properties of organic liquids. Journal of the American Chemical Society, 118(45), pp.11225–11236.

91. Abele, R. and Tampé, R., 2004. The ABCs of immunology: structure and function of TAP, the transporter associated with antigen processing. Physiology, 19(4), pp.216–224.

92. Nagpal, G., Gupta, S., Chaudhary, K., Dhanda, S.K., Prakash, S. and Raghava, G.P., 2015. VaccineDA: Prediction, design and genome-wide screening of oligodeoxynucleotide-based vaccine adjuvants. Scientific reports, 5, p.12478.

93. Dhanda, S. K., Vir, P. & Raghava, G. P. Designing of interferon-gamma inducing MHC class-II binders. Biol. Direct. 8, 30 (2013).

94. Saha, S. & Raghava, G. AlgPred: prediction of allergenic proteins and mapping of IgE epitopes. Nucleic. Acids. Res. 34, W202–W209 (2006).

95. Irini A Doytchinova and Darren R Flower. VaxiJen: a server for prediction of protective antigens, tumour antigens and subunit vaccines. BMC Bioinformatics. 2007 8:4.

96. Gasteiger, E., Hoogland, C., Gattiker, A., Duvaud, S.E., Wilkins, M.R., Appel, R.D. and Bairoch, A., 2005. Protein identification and analysis tools on the ExPASy server (pp. 571–607). Humana Press.

97. Morten Källberg, Haipeng Wang, Sheng Wang, Jian Peng, Zhiyong Wang, Hui Lu, and Jinbo Xu. Template-based protein structure modeling using the RaptorX web server. Nature Protocols 7, 1511–1522, 2012.

98. Ma, J., Wang, S., Zhao, F. and Xu, J., 2013. Protein threading using context-specific alignment potential. Bioinformatics, 29(13), pp.i257–i265.

99. Wang, Z., Zhao, F., Peng, J. and Xu, J., 2010, December. Protein 8-class secondary structure prediction using conditional neural fields. In 2010 IEEE International Conference on Bioinformatics and Biomedicine (BIBM) (pp. 109–114). IEEE.

100. Wang, Z. and Xu, J., 2013. Predicting protein contact map using evolutionary and physical constraints by integer programming. Bioinformatics, 29(13), pp.i266–i273.

101. Dong Xu and Yang Zhang. Improving the Physical Realism and Structural Accuracy of Protein Models by a Two-step Atomic-level Energy Minimization. Biophysical Journal, vol 101, 2525–2534 (2011).

102. J. Ko, H. Park, L. Heo, and C. Seok, GalaxyWEB server for protein structure prediction and refinement, Nucleic Acids Res. 40 (W1), W294–W297 (2012).

103. Shin, W.H., Lee, G.R., Heo, L., Lee, H. and Seok, C., 2014. Prediction of protein structure and interaction by GALAXY protein modeling programs. Bio Design, 2(1), pp.1–11.

104. Ramakrishnan, C. and Ramachandran, G.N., 1965. Stereochemical criteria for polypeptide and protein chain conformations: II. Allowed conformations for a pair of peptide units. Biophysical journal, 5(6), pp.909–933.

105. S.C. Lovell, I.W. Davis, W.B. Arendall III, P.I.W. de Bakker, J.M. Word, M.G. Prisant, J.S. Richardson and D.C. Richardson (2002) Structure validation by Calpha geometry: phi, psi and Cbeta deviation. Proteins: Structure, Function & Genetics. 50: 437–450.

106. Bell, J.K., Botos, I., Hall, P.R., Askins, J., Shiloach, J., Segal, D.M. and Davies, D.R., 2005. The molecular structure of the Toll-like receptor 3 ligand-binding domain. Proceedings of the National Academy of Sciences of the United States of America, 102(31), pp.10976–10980.

107. Duhovny D, Nussinov R, Wolfson HJ. Efficient Unbound Docking of Rigid Molecules. In Gusfield, et al., Ed. Proceedings of the 2’nd Workshop on Algorithms in Bioinformatics(WABI) Rome, Italy, Lecture Notes in Computer Science 2452, pp. 185–200, Springer Verlag, 2002

108. Schneidman-Duhovny D, Inbar Y, Nussinov R, Wolfson HJ. PatchDock and SymmDock: servers for rigid and symmetric docking. Nucl. Acids. Res. 33: W363–367, 2005.

109. Krieger, E. and Vriend, G., 2015. New ways to boost molecular dynamics simulations.Journal of computational chemistry, 36(13), pp.996–1007.

110. Maier, J.A., Martinez, C., Kasavajhala, K., Wickstrom, L., Hauser, K.E. and Simmerling, C., 2015. ff14SB: improving the accuracy of protein side chain and backbone parameters from ff99SB. Journal of chemical theory and computation, 11(8), pp.3696–3713.

111. Toukmaji, A., Sagui, C., Board, J. and Darden, T., 2000. Efficient particle-mesh Ewald based approach to fixed and induced dipolar interactions. The Journal of chemical physics, 113(24), pp.10913–10927.

112. Nezafat, N., Eslami, M., Negahdaripour, M., Rahbar, M.R. and Ghasemi, Y., 2017. Designing an efficient multi-epitope oral vaccine against Helicobacter pylori using immunoinformatics and structural vaccinology approaches. Molecular BioSystems, 13(4), pp.699–713.

113. Morla, S., Makhija, A. & Kumar, S. Synonymous codon usage pattern in glycoprotein gene of rabies virus. Gene. 584, 1–6 (2016).

114. Wu, X., Wu, S., Li, D., Zhang, J., Hou, L., Ma, J., Liu, W., Ren, D., Zhu, Y. and He, F., 2010. Computational identification of rare codons of Escherichia coli based on codon pairs preference. Bmc Bioinformatics, 11(1), p.61.

