## Supplementary v2 for "Exploring the structural basis to develop efficient multi-epitope vaccines displaying interaction with HLA and TAP and TLR3 molecules to prevent NIPAH infection, a global threat to human health"

**Supplementary table S1. Protein sequence retrieval, tertiary structures retrieval and homology modeling of nine Nipah proteins.** Nipah protein sequences were retrieved from NCBI. Available structure files (pdb) for Nipah proteins were retrieved from RCSB PDB. Nipah proteins with no tertiary structure available were subjected to homology modeling by Swissmodel.

| S.No | NIPAH-CoV Protein | Number of sequence retrieved from NCBI/PDB | PDB ID of available structure | Template used for modeling | QMEAN |
|---|---|---|---|---|---|
| 1 | C Protein | 13 | - | 5hyb.1.B | - 2.27 |
| 2 | Fusion Protein | 18 | 5EVM | - | - |
| 3 | Glycoprotein | 21 | 2VWD | - | - |
| 4 | Matrix Protein | 18 | - | 6bk6.1.A | - 1.81 |
| 5 | Nucleocapsid | 36 | 4CO6 | - | - |
| 6 | Phosphoprotein | 18 | 4N5B | - | - |
| 7 | Polymerase | 18 | - | 5a22.1.A | -6.06 |
| 8 | V protein | 11 | - | 4co6.1.B | -0.11 |
| 9 | W protein | 08 | 6BW0 | - | - |

**Supplementary table S2. Homology modeling for HLA alleles.** Tertiary structure of HLA alleles were modeled by homology modeling using SwissModel server. Templates were chosen with highest sequence identity. Generated models with acceptable QMEAN value were chosen for further studies.

| S.No | HLA class I allele | Template used for modeling | % sequence identity | QMEAN |
|---|---|---|---|---|
| 1 | A0101 | 4nqx.2.A | 100% | (-)0.07 |
| 2 | A3002 | 6eny.1.D | 96.48% | (-)0.52 |
| 3 | A2301 | 2bck.1.A | 98.91% | (-)0.01 |
| 4 | A2402 | 2bck.1.A | 100% | (-)0.02 |
| 5 | B1501 | 5txs.1.A | 100% | 0.14 |
| 6 | B3501 | 1a9b.1.A | 100% | (-)0.60 |
| 7 | A1101 | 6eny.1.D | 97.95% | 0.75 |
| 8 | A3201 | 6ei2.1.A | 92.00% | 0.57 |
| 9 | B4402 | 1m6o.1.A | 100% | 0.75 |
| 10 | B4403 | 4jqx.1.A | 100% | 0.64 |
| S.No | HLA class II allele | Template used for modeling | % sequence identity | QMEAN |
| 2 | DPA1-0103 | 4p4r.1.A | 100% | (-)0.73 |
| 2 | DRB3-0101 | 2q6w.1.B | 100% | (-)0.65 |
| 6 | DPB1-0201 | 4p5m.1.B | 98.42% | (-)0.37 |
| 4 | DRB1-0301 | 1a6a.1.B | 100% | (-)1.65 |

**Supplementary table S3. Shortlisted high scoring CTL epitopes.** Selected high scoring CTL epitopes and their respective HLA alleles binders are listed. *In-silico* analysis have shown all the selected epitopes to be non-toxic (Non-Toxin) as well as they show significant conservancy. ToxinPred analysis is based on the ToxinPred main dataset used by "ToxinPred" algorithm to predict toxicity of any unknown peptide. # Epitope match with previous studies indicating consensus in epitope screening by different approaches and methods.

| S.No | NIPAH Proteins | Epitope | Epitopes chosen for detailed study | Position | Length | HLA Class I Alleles | Percent of protein sequence matches at 100% identity (Conservancy) | ToxinPred Study |
|---|---|---|---|---|---|---|---|---|
| 1 | C Protein | MMASILLTLF | | 1-10 | 10 | B*15:01 | 100.00% (60/60) | Non-Toxin |
| 2 | Fusion Protein | AQITAGVALY | | 126-135 | 10 | B*15:01 | 97.30% (36/37) | Non-Toxin |
| 3 | Fusion Protein | KYLSDLLFVF | √ | 205-214 | 10 | A*23:01<br>A*24:02 | 97.30% (36/37) | Non-Toxin |
| 4 | Fusion Protein | MTIQAISQAF | | 226-235 | 10 | B*15:01 | 97.30% (36/37) | Non-Toxin |
| 5 | Fusion Protein | FALSNGVLF# | | 376-384 | 9 | B*35:01 | 100.00% (37/37) | Non-Toxin |
| 6 | Glycoprotein | LAMDEGYFAY | | 222-231 | 10 | B*35:01 | 90.91% (60/66) | Non-Toxin |
| 7 | Glycoprotein | TVYHCSAVY# | | 278-286 | 9 | A*30:02 | 100.00% (66/66) | Non-Toxin |
| 8 | Glycoprotein | AVYNNEFYY | √ | 284-292 | 9 | A*11:01<br>A*30:02 | 90.91% (60/66) | Non-Toxin |
| 9 | Glycoprotein | AENPVFTVF | √ | 532-540 | 9 | B*44:03<br>B*44:02 | 100.00% (66/66) | Non-Toxin |
| 10 | Matrix Protein | NYMYLICYGF | | 61-70 | 10 | A*23:01 | 97.44% (38/39) | Non-Toxin |
| 11 | Matrix Protein | YMIPRTMLEF | | 187-196 | 10 | B*15:01 | 97.44% (38/39) | Non-Toxin |
| 12 | Nucleocapsid | TPFVDSRAY | | 143-151 | 9 | B*35:01 | 100.00% (13/13) | Non-Toxin |
| 13 | Nucleocapsid | EIISDIGNY | | 250-258 | 9 | A*26:01 | 100.00% (13/13) | Non-Toxin |
| 14 | Nucleocapsid | YPALALNEF# | | 279-287 | 9 | B*35:01 | 100.00% (13/13) | Non-Toxin |
| 15 | Phosphoprotein/V Protein/ W protein | LDPVVTDVVY | √ | 107-116 | 10 | B*35:01 | 100.00% (27/27) | Non-Toxin |
| 16 | Phosphoprotein/V Protein/ W protein | LVSDAKMLSY | | 150-159 | 10 | A*01:01 | 59.26% (16/27) | Non-Toxin |
| 17 | Phosphoprotein/V Protein/ W protein | VSDAKMLSY# | √ | 151-159 | 9 | A*01:01 | 59.26% (16/27) | Non-Toxin |
| 18 | Phosphoprotein | MPSDDFSNTF | | 479-488 | 10 | B*35:01 | 96.30% (26/27) | Non-Toxin |
| 19 | Polymerase | YPECNNILF# | | 88-96 | 9 | B*35:01 | 90.00% (9/10) | Non-Toxin |
| 20 | Polymerase | IMKKSFKAY | | 110-118 | 9 | B*15:01 | 100.00% (10/10) | Non-Toxin |
| 21 | Polymerase | KWYECFLFWF | | 165-174 | 10 | A*23:01 | 100.00% (10/10) | Non-Toxin |
| 22 | Polymerase | FPVMGNRIY# | | 276-284 | 9 | B*35:01 | 100.00% (10/10) | Non-Toxin |
| 23 | Polymerase | AEFFSFFRTF# | √ | 354-363 | 10 | B*44:03 | 100.00% (10/10) | Non-Toxin |
| 24 | Polymerase | IPFLFLSAY# | | 811-819 | 9 | B*35:01 | 100.00% (10/10) | Non-Toxin |
| 25 | Polymerase | IATVYYTWAY | | 1300-1308 | 9 | B*35:01 | 100.00% (10/10) | Non-Toxin |
| 26 | Polymerase | LETDDYNGIY | | 1411-1420 | 10 | A*01:01 | 100.00% (10/10) | Non-Toxin |
| 27 | Polymerase | ETDDYNGIY# | | 1412-1420 | 9 | A*01:01 | 100.00% (10/10) | Non-Toxin |
| 28 | Polymerase | SQNLLVTSY# | | 1624-1632 | 9 | B*15:01 | 100.00% (10/10) | Non-Toxin |
| 29 | Polymerase | TSDLDFVIFY | | 1716-1725 | 10 | A*01:01 | 100.00% (10/10) | Non-Toxin |
| 30 | Polymerase | FPISRLFNMY | | 1983-1992 | 10 | B*35:01 | 100.00% (10/10) | Non-Toxin |
| 31 | Polymerase | RLFNMYRSY | √ | 1987-1995 | 9 | A*32:01<br>B*15:01 A*30:02 | 100.00% (10/10) | Non-Toxin |
| 32 | Polymerase | SYFGLVLVCF | | 1994-2003 | 10 | A*23:01 | 100.00% (10/10) | Non-Toxin |
| 33 | Polymerase | KYYQIDQPFF | | 2068-2077 | 10 | A*23:01 | 100.00% (10/10) | Non-Toxin |

**Supplementary table S4. Shortlisted high scoring HTL epitopes.** Selected high scoring HTL epitopes and their respective HLA alleles binders are listed above. *In-silico* analysis have shown all the selected epitopes to be non-toxic (Non-Toxin) as well as they show significant conservancy.

| S.No | NIPAH Protein | Epitope | Epitopes chosen for detailed study | Position | Length | HLA Class II Alleles | Percent of protein sequence matches at 100% identity (Conservancy) | ToxinPred Study |
|---|---|---|---|---|---|---|---|---|
| 1 | C Protein | VQMTYNWTQWLQTLY | | 103-117 | 15 | DPA1*01:03 DPB1*02:01 | 95.00% (57/60) | Non-Toxin |
| 2 | Fusion protein | DLALSKYLSDLLFVF | √ | 104-118 | 15 | DRB3*01:01 | 97.30% (36/37) | Non-Toxin |
| 3 | Fusion protein | LALSKYLSDLLFVFG | √ | 59-73 | 15 | DRB3*01:01 | 97.30% (36/37) | Non-Toxin |
| 4 | Fusion protein | ALSKYLSDLLFVFGP | √ | 99-113 | 15 | DRB3*01:01 | 97.30% (36/37) | Non-Toxin |
| 5 | Fusion protein | LSKYLSDLLFVFGPN | √ | 100-114 | 15 | DRB3*01:01 | 97.30% (36/37) | Non-Toxin |
| 6 | Fusion protein | SKYLSDLLFVFGPNL | √ | 101-115 | 15 | DRB3*01:01 | 97.30% (36/37) | Non-Toxin |
| 7 | Fusion protein | KYLSDLLFVFGPNLQ | √ | 102-116 | 15 | DRB3*01:01 | 97.30% (36/37) | Non-Toxin |
| 8 | Fusion protein | YLSDLLFVFGPNLQD | √ | 103-117 | 15 | DRB3*01:01 | 97.30% (36/37) | Non-Toxin |
| 9 | Glycoprotein | ASFSWDTMIKFGDVL | | 456-470 | 15 | DRB3*01:01 | 19.70% (13/66) | Non-Toxin |
| 10 | Glycoprotein | FSWDTMIKFGDVLTV | | 458-472 | 15 | DRB3*01:01 | 19.70% (13/66) | Non-Toxin |
| 11 | Glycoprotein | GVYNDAFLIDRINWI | | 506-520 | 15 | DRB3*01:01 | 100.00% (66/66) | Non-Toxin |
| 12 | Glycoprotein | NDAFLIDRINWISAG | | 509-523 | 15 | DRB3*01:01 | 100.00% (66/66) | Non-Toxin |
| 13 | Glycoprotein | DAFLIDRINWISAGV | | 510-424 | 15 | DRB3*01:01 | 100.00% (66/66) | Non-Toxin |
| 14 | Glycoprotein | AFLIDRINWISAGVF | | 511-525 | 15 | DRB3*01:01 | 100.00% (66/66) | Non-Toxin |
| 15 | Glycoprotein | FLIDRINWISAGVFL | | 512-526 | 15 | DRB3*01:01 | 100.00% (66/66) | Non-Toxin |
| 16 | Matrix Protein | IPREFMIYDDVFIDN | | 331-345 | 15 | DRB3*01:01 | 38.46% (15/39) | Non-Toxin |
| 17 | Matrix Protein | FMIYDDVFIDNTGRI | | 335-349 | 15 | DRB3*01:01 | 94.87% (37/39) | Non-Toxin |
| 18 | Nucleocapsid | LSSDQVAELAAAVQE | | 384-398 | 15 | DQA1*04:01 DQB1*04:02 | 47.37% (9/19) | Non-Toxin |
| 19 | Nucleocapsid | SSDQVAELAAAVQET | | 385-399 | 15 | DQA1*04:01 DQB1*04:02 | 47.37% (9/19) | Non-Toxin |
| 20 | Nucleocapsid | SDQVAELAAAVQETS | | 386-400 | 15 | DQA1*04:01 DQB1*04:02 | 47.37% (9/19) | Non-Toxin |
| 21 | Nucleocapsid | DQVAELAAAVQETSA | | 387-401 | 15 | DQA1*04:01 DQB1*04:02 | 47.37% (9/19) | Non-Toxin |
| 22 | Nucleocapsid | QVAELAAAVQETSAG | | 388-402 | 15 | DQA1*04:01 DQB1*04:02 | 100.00% (19/19) | Non-Toxin |
| 23 | Phosphoprotein/V Protein/ W protein | NNGNVCLVSDAKMLS | √ | 144-158 | 15 | DRB1*03:01 | 62.96% (17/27) | Non-Toxin |
| 24 | Phosphoprotein/V Protein/ W protein | NGNVCLVSDAKMLSY | √ | 145-159 | 15 | DRB1*03:01 | 59.26% (16/27) | Non-Toxin |
| 25 | Phosphoprotein/V Protein/ W protein | GNVCLVSDAKMLSYA | √ | 146-160 | 15 | DRB1*03:01 | 59.26% (16/27) | Non-Toxin |
| 26 | Phosphoprotein/V Protein/ W protein | NVCLVSDAKMLSYAP | √ | 147-161 | 15 | DRB1*03:01 | 59.26% (16/27) | Non-Toxin |
| 27 | Phosphoprotein/V Protein/ W protein | VCLVSDAKMLSYAPE | √ | 148-162 | 15 | DRB1*03:01 | 59.26% (16/27) | Non-Toxin |
| 28 | Polymerase | NIDNIHLLAEFFSFF | √ | 346-360 | 15 | DPA1*01:03 DPB1*02:01 | 100.00% (10/10) | Non-Toxin |
| 29 | Polymerase | IDNIHLLAEFFSFFR | √ | 347-361 | 15 | DPA1*01:03 DPB1*02:01 | 100.00% (10/10) | Non-Toxin |
| 30 | Polymerase | DNIHLLAEFFSFFRT | √ | 348-362 | 15 | DPA1*01:03 DPB1*02:01 | 100.00% (10/10) | Non-Toxin |
| 31 | Polymerase | NIHLLAEFFSFFRTF | √ | 349-363 | 15 | DPA1*01:03 DPB1*02:01 | 100.00% (10/10) | Non-Toxin |
| 32 | Polymerase | IHLLAEFFSFFRTFG | √ | 350-364 | 15 | DPA1*01:03 DPB1*02:01 | 100.00% (10/10) | Non-Toxin |
| 33 | Polymerase | LELASFLMDRRVILP | | 1102-1116 | 15 | DRB3*01:01 | 100.00% (10/10) | Non-Toxin |
| 34 | Polymerase | ELASFLMDRRVILPR | | 1103-1117 | 15 | DRB3*01:01 | 100.00% (10/10) | Non-Toxin |
| 35 | Polymerase | LASFLMDRRVILPRA | | 1104-1118 | 15 | DRB3*01:01 | 100.00% (10/10) | Non-Toxin |
| 36 | Polymerase | ASFLMDRRVILPRAA | | 1105-1119 | 15 | DRB3*01:01 | 100.00% (10/10) | Non-Toxin |
| 37 | Polymerase | LDFVIFYASLTYLRR | | 1719-1733 | 15 | DPA1*02:01 DPB1*14:01 | 100.00% (10/10) | Non-Toxin |
| 38 | Polymerase | FVIFYASLTYLRRGI | | 1721-1735 | 15 | DPA1*02:01 DPB1*14:01 | 100.00% (10/10) | Non-Toxin |

**Supplementary table S5. World population coverage by the shortlisted CTL and HTL epitopes combined.** The cumulative percent of world (countries as listed in table) population coverage is 97.88% by the joint administration of selected CTL and HTL epitopes as vaccine candidate.

a projected population coverage
b average number of epitope hits / HLA combinations recognized by the population
c minimum number of epitope hits / HLA combinations recognized by 90% of the population

| Population/Area | Class combined | | |
|---|---|---|---|
| | coverage[a] | average_hit[b] | pc90[c] |
| Austria | 86.74% | 7.22 | 0.75 |
| Belgium | 82.97% | 5.68 | 0.59 |
| Borneo | 9.26% | 0.52 | 0.55 |
| Brazil | 99.93% | 13.73 | 7.16 |
| Canada | 27.97% | 1.53 | 0.69 |
| Central Africa | 95.73% | 10.45 | 2.84 |
| Central America | 50.90% | 2.48 | 0.41 |
| Chile | 81.43% | 4.35 | 0.54 |
| China | 96.24% | 8.75 | 2 |
| Cuba | 81.77% | 5.29 | 0.55 |
| Czech Republic | 82.95% | 7.12 | 0.59 |
| Denmark | 27.87% | 1.51 | 0.69 |
| East Africa | 95.36% | 10.37 | 2.87 |
| East Asia | 93.70% | 7.99 | 1.29 |
| England | 90.31% | 7.45 | 1.03 |
| Europe | 99.91% | 13.69 | 6.97 |
| Finland | 85.91% | 8.29 | 0.71 |
| France | 99.98% | 13.86 | 7.26 |
| Georgia | 88.89% | 7.81 | 0.9 |
| Germany | 89.61% | 7.97 | 0.96 |
| Hong Kong | 75.06% | 1.63 | 0.4 |
| India | 99.13% | 10.73 | 6.15 |
| Indonesia | 70.81% | 2.13 | 0.34 |
| Iran | 74.68% | 4.17 | 0.39 |
| Ireland South | 91.40% | 7.24 | 1.14 |
| Israel | 72.73% | 3.85 | 0.37 |
| Italy | 95.31% | 10.89 | 2.76 |
| Japan | 98.13% | 11.21 | 3.28 |
| Jordan | 55.10% | 1.58 | 0.22 |
| Kenya | 94.26% | 9.77 | 2.59 |
| Korea; South | 86.85% | 5.82 | 0.76 |
| Lebanon | 54.37% | 3.41 | 0.44 |
| Malaysia | 66.85% | 2.91 | 0.3 |
| Mexico | 99.99% | 13.79 | 6.69 |
| Mongolia | 87.73% | 5.75 | 0.82 |
| Netherlands | 45.52% | 2.86 | 0.37 |
| North Africa | 85.76% | 7.63 | 0.7 |
| North America | 99.99% | 13.61 | 6.86 |
| Northeast Asia | 96.28% | 8.71 | 1.98 |
| Norway | 49.89% | 3.23 | 0.4 |
| Oceania | 97.70% | 9.04 | 2.13 |
| Oman | 81.20% | 3.52 | 0.53 |
| Pakistan | 71.59% | 1.54 | 0.35 |
| Peru | 83.62% | 6.66 | 0.61 |
| Philippines | 63.66% | 0.79 | 0.28 |

| Population/Area | Class combined | | |
|---|---|---|---|
| | coverage[a] | average_hit[b] | pc90[c] |
| Poland | 83.36% | 5.59 | 0.6 |
| Portugal | 81.92% | 5.06 | 0.55 |
| Romania | 80.31% | 4.61 | 0.51 |
| Russia | 99.80% | 14.12 | 7.27 |
| Saudi Arabia | 73.55% | 4.44 | 0.38 |
| Scotland | 55.20% | 1.76 | 0.22 |
| Singapore | 75.29% | 2.61 | 0.4 |
| Slovakia | 45.98% | 2.94 | 0.37 |
| South Africa | 75.52% | 4.38 | 0.41 |
| South America | 97.92% | 10.72 | 4.7 |
| South Asia | 99.33% | 10.98 | 6.22 |
| Southeast Asia | 82.34% | 3.19 | 0.57 |
| Southwest Asia | 79.01% | 5.18 | 0.48 |
| Spain | 99.93% | 10.55 | 6.29 |
| Sri Lanka | 38.06% | 1.23 | 0.16 |
| Sri Lanka Asian | 38.06% | 1.23 | 0.16 |
| Sweden | 99.96% | 15.31 | 7.31 |
| Taiwan | 90.12% | 4.26 | 1 |
| Thailand | 78.15% | 3.4 | 0.46 |
| Turkey | 23.60% | 1.25 | 0.65 |
| United Arab Emirates | 33.67% | 1.73 | 0.71 |
| United Kingdom | 46.56% | 1.86 | 0.37 |
| United States | 100.00% | 13.38 | 6.81 |
| Vietnam | 75.66% | 2.48 | 0.41 |
| West Africa | 98.14% | 11.01 | 2.78 |
| West Indies | 85.36% | 6.22 | 0.68 |
| Zimbabwe | 82.41% | 6.54 | 0.57 |
| **Cumulative percent of world population coverage** | **97.88%** | **11.33** | **4.99** |
| **Average population coverage** | **77.84** | **6.38** | **1.87** |
| **Standard deviation** | **21.97** | **4.07** | **2.3** |

**Supplementary table S6. Shortlisted B Cell epitopes.** BepiPred Linear B Cell epitopes showing sequence overlap with CTL and HTL epitopes are shortlisted above. *In-silico* analysis have shown all the selected epitopes to be non-toxic (Non-Toxin) as well as they show significant amino acid sequence conservancy. #Epitope match with previous studies indicating consensus in epitope screening by different approaches and methods.

| S.No. | NIPAH Protein | B Cell Epitope | Position | Length | Percent of protein sequence matches at 100% identity (Conservancy) | ToxinPred Study |
|---|---|---|---|---|---|---|
| 1 | Fusion Protein | GPNLQDPVSNSM | 215-226 | 12 | 97.30% (36/37) | Non-Toxin |
| 2 | Glycoprotein | WTPPNPNT | 271-278 | 8 | 22.73% (15/66) | Non-Toxin |
| 3 | Glycoprotein | SWDTMI | 459-464 | 6 | 100.00% (66/66) | Non-Toxin |
| 4 | Glycoprotein | NQTAE | 529-533 | 5 | 100.00% (66/66) | Non-Toxin |
| 5 | Matrix Protein | SGIYM | 184-188 | 5 | 97.44% (38/39) | Non-Toxin |
| 6 | Matrix Protein | SIPREFMIY# | 330-338 | 9 | 38.46% (15/39) | Non-Toxin |
| 7 | Matrix Protein | DVFIDNTGRI | 340-349 | 10 | 94.87% (37/39) | Non-Toxin |
| 8 | Phosphoprotein | GYGFTSSPERGWSDYTSGA | 125-143 | 19 | 51.85% (14/27) | Non-Toxin |
| 9 | Phosphoprotein | IAVSKEDR | 163-170 | 8 | 100.00% (27/27) | Non-Toxin |
| 10 | Polymerase | NIDN | 346-349 | 4 | 100.00% (10/10) | Non-Toxin |
| 11 | Polymerase | GHPILE | 364-369 | 6 | 100.00% (10/10) | Non-Toxin |
| 12 | Polymerase | DKSFDEDLEL | 1095-1104 | 10 | 100.00% (10/10) | Non-Toxin |
| 13 | Polymerase | LMDR | 1108-1111 | 4 | 100.00% (10/10) | Non-Toxin |
| 14 | Polymerase | LRLETDDYNG | 1409-1418 | 10 | 100.00% (10/10) | Non-Toxin |
| 15 | Polymerase | GFPIS | 1982-1986 | 5 | 100.00% (10/10) | Non-Toxin |
| 16 | Polymerase | PVYSNPD | 2004-2010 | 7 | 100.00% (10/10) | Non-Toxin |

**Supplementary table S7. CTL epitope prediction.** Detailed scoring of all screened CTL epitopes and their respective HLA class I allele binders. CTL epitopes were chosen on the basis of high “Total score” and higher number of HLA allele binders. Total score is a combined score of TAP score, MHC score, Proteasome score and Processing score.

| S.No. | NIPAH-CoV Protein | Epitope | HLA Class I Alleles | Proteasome Score | TAP Score | MHC Score | Processing Score | Total Score | MHC IC50[nM] | Immunogenicity-Score |
|---|---|---|---|---|---|---|---|---|---|---|
| 1 | C Protein | MMASILLTLF | B*15:01 | 1.2 | 1.13 | -0.76 | 2.33 | 1.57 | 5.8 | -0.01913 |
| 2 | Fusion Protein | AQITAGVALY | B*15:01 | 1.18 | 1.46 | -1 | 2.64 | 1.64 | 10 | 0.21876 |
| 3 | Fusion Protein | KYLSDLLFVF | A*23:01 | 1.55 | 1.29 | -0.74 | 2.84 | 2.1 | 5.5 | -0.04679 |
| | Fusion Protein | | A*24:02 | 1.55 | 1.29 | -0.98 | 2.84 | 1.86 | 9.5 | -0.04679 |
| 4 | Fusion Protein | MTIQAISQAF | B*15:01 | 1.4 | 1.16 | -0.92 | 2.57 | 1.64 | 8.4 | -0.13629 |
| 5 | Fusion Protein | FALSNGVLF | B*35:01 | 1.49 | 1.15 | -0.77 | 2.64 | 1.86 | 5.9 | -0.11611 |
| 6 | Glycoprotein | LAMDEGYFAY | B*35:01 | 1.43 | 1.34 | -0.85 | 2.77 | 1.93 | 7 | 0.214 |
| 7 | Glycoprotein | TVYHCSAVY | A*30:02 | 1.57 | 1.46 | -1.51 | 3.03 | 1.52 | 32.6 | -0.11974 |
| 8 | Glycoprotein | AVYNNEFYY | A*11:01 | 1.21 | 1.5 | -1.07 | 2.7 | 1.63 | 11.8 | 0.17688 |
| | Glycoprotein | | A*30:02 | 1.21 | 1.5 | -1.11 | 2.7 | 1.59 | 13 | 0.17688 |
| 9 | Glycoprotein | AENPVFTVF | B*44:03 | 1.71 | 1.07 | -1.09 | 2.77 | 1.69 | 12.3 | 0.19402 |
| | Glycoprotein | | B*44:02 | 1.71 | 1.07 | -1.21 | 2.77 | 1.56 | 16.3 | 0.19402 |
| 10 | Matrix Protein | NYMYLICYGF | A*23:01 | 1.29 | 1.32 | -1.09 | 2.61 | 1.52 | 12.2 | 0.02401 |
| 11 | Matrix Protein | YMIPRTMLEF | B*15:01 | 1.4 | 1.18 | -0.79 | 2.57 | 1.79 | 6.1 | 0.00408 |
| 12 | Nucleocapsid | TPFVDSRAY | B*35:01 | 1.33 | 1.22 | -0.79 | 2.56 | 1.77 | 6.1 | 0.01195 |
| 13 | Nucleocapsid | EIISDIGNY | A*26:01 | 1.19 | 1.29 | -0.6 | 2.47 | 1.87 | 4 | 0.04843 |
| 14 | Nucleocapsid | YPALALNEF | B*35:01 | 1.25 | 1.02 | -0.61 | 2.27 | 1.66 | 4.1 | 0.08224 |
| 15 | Phosphoprotein/V Protein/ W protein | LDPVVTDVVY | B*35:01 | 1.74 | 1.11 | -1.34 | 2.85 | 1.51 | 22.1 | 0.19578 |
| 16 | Phosphoprotein/V Protein/ W protein | LVSDAKMLSY | A*01:01 | 1.48 | 1.33 | -1.26 | 2.81 | 1.55 | 18 | -0.4746 |
| 17 | Phosphoprotein/V Protein/ W protein | VSDAKMLSY | A*01:01 | 1.48 | 1.26 | -0.94 | 2.74 | 1.8 | 8.7 | -0.43475 |
| 18 | Phosphoprotein | MPSDDFSNTF | B*35:01 | 1.53 | 0.99 | -0.88 | 2.52 | 1.64 | 7.6 | -0.03429 |
| 19 | Polymerase | YPECNNILF | B*35:01 | 1.47 | 0.96 | -0.82 | 2.44 | 1.62 | 6.6 | 0.0717 |
| 20 | Polymerase | IMKKSFKAY | B*15:01 | 1.59 | 1.39 | -1.07 | 2.98 | 1.92 | 11.7 | -0.49704 |
| 21 | Polymerase | KWYECFLFWF | A*23:01 | 1.24 | 1.36 | -1 | 2.6 | 1.6 | 10.1 | 0.37883 |
| 22 | Polymerase | FPVMGNRIY | B*35:01 | 1.57 | 1.15 | -0.51 | 2.72 | 2.22 | 3.2 | -0.01495 |
| 13 | Polymerase | AEFFSFFRTF | B*44:03 | 1.49 | 1.17 | -1.05 | 2.66 | 1.61 | 11.3 | 0.28526 |
| 24 | Polymerase | IPFLFLSAY | B*35:01 | 1.39 | 1.2 | -0.67 | 2.59 | 1.92 | 4.7 | 0.01364 |
| 25 | Polymerase | IATVYTWAY | B*35:01 | 1.56 | 1.34 | -0.93 | 2.9 | 1.97 | 8.6 | 0.29688 |
| 26 | Polymerase | LETDDYNGIY | A*01:01 | 1.41 | 1.25 | -0.93 | 2.66 | 1.73 | 8.6 | 0.15319 |
| 27 | Polymerase | ETDDYNGIY | A*01:01 | 1.41 | 1.12 | -0.91 | 2.53 | 1.62 | 8.1 | 0.12619 |
| 28 | Polymerase | SQNLLVTSY | B*15:01 | 1.6 | 1.31 | -1.38 | 2.9 | 1.52 | 24.2 | -0.0491 |
| 29 | Polymerase | TSDLDFVIFY | A*01:01 | 1.68 | 1.26 | -0.87 | 2.94 | 2.07 | 7.4 | 0.35122 |
| 30 | Polymerase | FPISRLFNMY | B*35:01 | 1.46 | 1.11 | -0.98 | 2.57 | 1.59 | 9.6 | -0.08153 |
| 31 | Polymerase | RLFNMYRSY | A*32:01 | 1.42 | 1.46 | -1.11 | 2.88 | 1.77 | 12.8 | -0.19597 |
| | Polymerase | | B*15:01 | 1.42 | 1.46 | -1.18 | 2.88 | 1.69 | 15.3 | -0.19597 |
| | Polymerase | | A*30:02 | 1.42 | 1.46 | -1.2 | 2.88 | 1.68 | 15.8 | -0.19597 |
| 32 | Polymerase | SYFGLVLVCF | A*23:01 | 1.48 | 1.34 | -1.04 | 2.82 | 1.78 | 10.9 | 0.0944 |
| 33 | Polymerase | KYYQIDQPFF | A*23:01 | 1.33 | 1.39 | -0.98 | 2.72 | 1.73 | 9.6 | -0.01656 |

**Supplementary table S8. HTL epitope prediction.** Percentile rank of HTL epitopes and their respective HLA class II allele binders. HTL epitopes were screened on the basis of percentile rank (lower the percentile number, higher the rank) and larger number of HLA allele binders. Last column show the method used for epitope screening.

| S.No | Epitope | Epitope | HLA Class II Alleles | Percentile rank | Method used |
|---|---|---|---|---|---|
| 1 | C Protein | VQMTYNWTQWLQTLY | DPA1*01:03 DPB1*02:01 | 0.08 | Consensus (comb.lib./smm/nn) |
| 2 | Fusion Protein | DLALSKYLSDLLFVF | DRB3*01:01 | 0.01 | Consensus (comb.lib./smm/nn) |
| 3 | Fusion Protein | LALSKYLSDLLFVFG | DRB3*01:01 | 0.01 | Consensus (comb.lib./smm/nn) |
| 4 | Fusion Protein | ALSKYLSDLLFVFGP | DRB3*01:01 | 0.01 | Consensus (comb.lib./smm/nn) |
| 5 | Fusion Protein | LSKYLSDLLFVFGPN | DRB3*01:01 | 0.01 | Consensus (comb.lib./smm/nn) |
| 6 | Fusion Protein | SKYLSDLLFVFGPNL | DRB3*01:01 | 0.01 | Consensus (comb.lib./smm/nn) |
| 7 | Fusion Protein | KYLSDLLFVFGPNLQ | DRB3*01:01 | 0.01 | Consensus (comb.lib./smm/nn) |
| 8 | Fusion Protein | YLSDLLFVFGPNLQD | DRB3*01:01 | 0.01 | Consensus (comb.lib./smm/nn) |
| 9 | Glycoprotein | ASFSWDTMIKFGDVL | DRB3*01:01 | 0.01 | Consensus (comb.lib./smm/nn) |
| 10 | Glycoprotein | FSWDTMIKFGDVLTV | DRB3*01:01 | 0.01 | Consensus (comb.lib./smm/nn) |
| 11 | Glycoprotein | GVYNDAFLIDRINWI | DRB3*01:01 | 0.01 | Consensus (comb.lib./smm/nn) |
| 12 | Glycoprotein | NDAFLIDRINWISAG | DRB3*01:01 | 0.01 | Consensus (comb.lib./smm/nn) |
| 13 | Glycoprotein | DAFLIDRINWISAGV | DRB3*01:01 | 0.01 | Consensus (comb.lib./smm/nn) |
| 14 | Glycoprotein | AFLIDRINWISAGVF | DRB3*01:01 | 0.01 | Consensus (comb.lib./smm/nn) |
| 15 | Glycoprotein | FLIDRINWISAGVFL | DRB3*01:01 | 0.01 | Consensus (comb.lib./smm/nn) |
| 16 | Matrix Protein | IPREFMIYDDVFIDN | DRB3*01:01 | 0.01 | Consensus (comb.lib./smm/nn) |
| 17 | Matrix Protein | FMIYDDVFIDNTGRI | DRB3*01:01 | 0.01 | Consensus (comb.lib./smm/nn) |
| 18 | Nucleocapsid | LSSDQVAELAAAVQE | DQA1*04:01 DQB1*04:02 | 0.01 | Consensus (comb.lib./smm/nn) |
| 19 | Nucleocapsid | SSDQVAELAAAVQET | DQA1*04:01 DQB1*04:02 | 0.01 | Consensus (comb.lib./smm/nn) |
| 20 | Nucleocapsid | SDQVAELAAAVQETS | DQA1*04:01 DQB1*04:02 | 0.01 | Consensus (comb.lib./smm/nn) |
| 21 | Nucleocapsid | DQVAELAAAVQETSA | DQA1*04:01 DQB1*04:02 | 0.01 | Consensus (comb.lib./smm/nn) |
| 22 | Nucleocapsid | QVAELAAAVQETSAG | DQA1*04:01 DQB1*04:02 | 0.01 | Consensus (comb.lib./smm/nn) |
| 23 | Phosphoprotein/V Protein/ W protein | NNGNVCLVSDAKMLS | DRB1*03:01 | 0.01 | Consensus (comb.lib./smm/nn) |
| 24 | Phosphoprotein/V Protein/ W protein | NGNVCLVSDAKMLSY | DRB1*03:01 | 0.01 | Consensus (comb.lib./smm/nn) |
| 25 | Phosphoprotein/V Protein/ W protein | GNVCLVSDAKMLSYA | DRB1*03:01 | 0.01 | Consensus (comb.lib./smm/nn) |
| 26 | Phosphoprotein/V Protein/ W protein | NVCLVSDAKMLSYAP | DRB1*03:01 | 0.01 | Consensus (comb.lib./smm/nn) |
| 27 | Phosphoprotein/V Protein/ W protein | VCLVSDAKMLSYAPE | DRB1*03:01 | 0.01 | Consensus (comb.lib./smm/nn) |
| 28 | Polymerase | NIDNIHLLAEFFSFF | DPA1*01:03 DPB1*02:01 | 0.01 | Consensus (comb.lib./smm/nn) |
| 29 | Polymerase | IDNIHLLAEFFSFFR | DPA1*01:03 DPB1*02:01 | 0.01 | Consensus (comb.lib./smm/nn) |
| 30 | Polymerase | DNIHLLAEFFSFFRT | DPA1*01:03 DPB1*02:01 | 0.01 | Consensus (comb.lib./smm/nn) |
| 31 | Polymerase | NIHLLAEFFSFFRTF | DPA1*01:03 DPB1*02:01 | 0.01 | Consensus (comb.lib./smm/nn) |
| 32 | Polymerase | IHLLAEFFSFFRTFG | DPA1*01:03 DPB1*02:01 | 0.01 | Consensus (comb.lib./smm/nn) |
| 33 | Polymerase | LELASFLMDRRVILP | DRB3*01:01 | 0.01 | Consensus (comb.lib./smm/nn) |
| 34 | Polymerase | ELASFLMDRRVILPR | DRB3*01:01 | 0.01 | Consensus (comb.lib./smm/nn) |
| 35 | Polymerase | LASFLMDRRVILPRA | DRB3*01:01 | 0.01 | Consensus (comb.lib./smm/nn) |
| 36 | Polymerase | ASFLMDRRVILPRAA | DRB3*01:01 | 0.01 | Consensus (comb.lib./smm/nn) |
| 37 | Polymerase | LDFVIFYASLTYLRR | DPA1*02:01 DPB1*14:01 | 0.01 | Consensus (comb.lib./smm/nn) |
| 38 | Polymerase | FVIFYASLTYLRRGI | DPA1*02:01 DPB1*14:01 | 0.01 | Consensus (comb.lib./smm/nn) |

**Supplementary table S9. INF-γ epitopes from CTL and HTL MEVs.** INF-γ inducing (POSITIVE) epitopes from CTL and HTL MEVs were screened by using "Motif and SVM hybrid" (MERCI & SVM) approaches.

| CLT Epitopes also predicted to be IFN-gamma epitopes | | | | | |
|---|---|---|---|---|---|
| S.No | Start-END | Sequence | Method | Result | Score |
| 1 | 22-37 | RRYKQIGTCGLPGTK | MERCI | POSITIVE | 1 |
| 2 | 23-38 | RYKQIGTCGLPGTKC | MERCI | POSITIVE | 1 |
| 3 | 24-39 | YKQIGTCGLPGTKCC | MERCI | POSITIVE | 1 |
| 4 | 25-40 | KQIGTCGLPGTKCCK | MERCI | POSITIVE | 1 |
| 5 | 26-41 | QIGTCGLPGTKCCKK | MERCI | POSITIVE | 1 |
| 6 | 38-53 | CKKPEAAAKMMASIL | MERCI | POSITIVE | 1 |
| 7 | 112-127 | GVLFGGGGSLAMDEG | MERCI | POSITIVE | 1 |
| 8 | 113-128 | VLFGGGGSLAMDEGY | MERCI | POSITIVE | 1 |
| 9 | 114-129 | LFGGGGSLAMDEGYF | MERCI | POSITIVE | 1 |
| 10 | 115-130 | FGGGGSLAMDEGYFA | MERCI | POSITIVE | 1 |
| 11 | 116-131 | GGGGSLAMDEGYFAY | MERCI | POSITIVE | 1 |
| 12 | 117-132 | GGGSLAMDEGYFAYG | MERCI | POSITIVE | 1 |
| 13 | 118-133 | GGSLAMDEGYFAYGG | MERCI | POSITIVE | 1 |
| 14 | 119-134 | GSLAMDEGYFAYGGG | MERCI | POSITIVE | 1 |
| 15 | 120-135 | SLAMDEGYFAYGGGG | MERCI | POSITIVE | 1 |
| 16 | 325-340 | KKSFKAYGGGGSKWY | MERCI | POSITIVE | 1 |
| 17 | 326-341 | KSFKAYGGGGSKWYE | MERCI | POSITIVE | 1 |
| 18 | 327-342 | SFKAYGGGGSKWYEC | MERCI | POSITIVE | 1 |
| 19 | 328-343 | FKAYGGGGSKWYECF | MERCI | POSITIVE | 1 |
| 20 | 329-344 | KAYGGGGSKWYECFL | MERCI | POSITIVE | 2 |
| 21 | 330-345 | AYGGGGSKWYECFLF | MERCI | POSITIVE | 2 |
| 22 | 331-346 | YGGGGSKWYECFLFW | MERCI | POSITIVE | 2 |
| 23 | 332-347 | GGGGSKWYECFLFWF | MERCI | POSITIVE | 2 |
| 24 | 333-348 | GGGSKWYECFLFWFG | MERCI | POSITIVE | 1 |
| 25 | 334-349 | GGSKWYECFLFWFGG | MERCI | POSITIVE | 1 |
| 26 | 335-350 | GSKWYECFLFWFGGG | MERCI | POSITIVE | 1 |
| 27 | 336-351 | SKWYECFLFWFGGGG | MERCI | POSITIVE | 1 |
| 28 | 337-352 | KWYECFLFWFGGGGS | MERCI | POSITIVE | 1 |
| 29 | 388-403 | AYGGGGSIATVYTWA | MERCI | POSITIVE | 1 |
| 30 | 392-407 | GGSIATVYTWAYGGG | MERCI | POSITIVE | 1 |
| 31 | 393-408 | GSIATVYTWAYGGGG | MERCI | POSITIVE | 1 |
| 32 | 394-409 | SIATVYTWAYGGGGS | MERCI | POSITIVE | 1 |
| 33 | 559-572 | STRGRKCCRRKKHHH | MERCI | POSITIVE | 1 |
| HLT Epitopes also predicted to be IFN-gamma epitopes | | | | | |
| S.No | Start-END | Sequence | Method | Result | Score |
| 1 | 22-37 | RRYKQIGTCGLPGTK | MERCI | POSITIVE | 1 |
| 2 | 23-38 | RYKQIGTCGLPGTKC | MERCI | POSITIVE | 1 |
| 3 | 24-39 | YKQIGTCGLPGTKCC | MERCI | POSITIVE | 1 |
| 4 | 25-40 | KQIGTCGLPGTKCCK | MERCI | POSITIVE | 1 |
| 5 | 26-41 | QIGTCGLPGTKCCKK | MERCI | POSITIVE | 1 |
| 6 | 38-53 | CKKPEAAAKVQMTYN | MERCI | POSITIVE | 1 |
| 7 | 379-394 | GRIGGGGSLSSDQVA | MERCI | POSITIVE | 1 |
| 8 | 380-395 | RIGGGGSLSSDQVAE | MERCI | POSITIVE | 1 |
| 9 | 381-396 | IGGGGSLSSDQVAEL | MERCI | POSITIVE | 1 |
| 10 | 382-397 | GGGGSLSSDQVAELA | MERCI | POSITIVE | 1 |
| 11 | 383-398 | GGGSLSSDQVAELAA | MERCI | POSITIVE | 1 |
| 12 | 482-496 | GGGGSNNGNVCLVSD | MERCI | POSITIVE | 1 |
| 13 | 483-497 | GGGSNNGNVCLVSDA | MERCI | POSITIVE | 1 |
| 14 | 484-498 | GGSNNGNVCLVSDAK | MERCI | POSITIVE | 1 |
| 15 | 485-499 | GSNNGNVCLVSDAKM | MERCI | POSITIVE | 1 |
| 16 | 486-500 | SNNGNVCLVSDAKML | MERCI | POSITIVE | 1 |
| 17 | 487-501 | NNGNVCLVSDAKMLS | MERCI | POSITIVE | 1 |
| 18 | 488-502 | NGNVCLVSDAKMLSG | MERCI | POSITIVE | 1 |
| 19 | 489-503 | GNVCLVSDAKMLSGG | MERCI | POSITIVE | 1 |
| 20 | 506-520 | SNGNVCLVSDAKMLS | MERCI | POSITIVE | 1 |
| 21 | 507-521 | NGNVCLVSDAKMLSY | MERCI | POSITIVE | 1 |
| 22 | 508-522 | GNVCLVSDAKMLSYG | MERCI | POSITIVE | 1 |
| 23 | 522-536 | GGGGSGNVCLVSDAK | MERCI | POSITIVE | 1 |
| 24 | 523-537 | GGGSGNVCLVSDAKM | MERCI | POSITIVE | 1 |

| S.No | Start-END | Sequence | Method | Result | Score |
|---|---|---|---|---|---|
| 25 | 524-538 | GGSGNVCLVSDAKML | MERCI | POSITIVE | 1 |
| 26 | 525-539 | GSGNVCLVSDAKMLS | MERCI | POSITIVE | 1 |
| 27 | 526-540 | SGNVCLVSDAKMLSY | MERCI | POSITIVE | 1 |
| 28 | 527-541 | GNVCLVSDAKMLSYA | MERCI | POSITIVE | 1 |
| 29 | 559-573 | YAPGGGGSVCLVSDA | MERCI | POSITIVE | 2 |
| 30 | 560-574 | APGGGGSVCLVSDAK | MERCI | POSITIVE | 2 |
| 31 | 561-575 | PGGGGSVCLVSDAKM | MERCI | POSITIVE | 2 |
| 32 | 562-576 | GGGGSVCLVSDAKML | MERCI | POSITIVE | 2 |
| 33 | 563-577 | GGGSVCLVSDAKMLS | MERCI | POSITIVE | 2 |
| 34 | 564-578 | GGSVCLVSDAKMLSY | MERCI | POSITIVE | 2 |
| 35 | 565-579 | GSVCLVSDAKMLSYA | MERCI | POSITIVE | 1 |
| 36 | 592-606 | HLLAEFFSFFGGGGS | MERCI | POSITIVE | 1 |
| 37 | 593-607 | LLAEFFSFFGGGGSI | MERCI | POSITIVE | 1 |
| 38 | 771-785 | IFYASLTYLRRGGGG | MERCI | POSITIVE | 7 |
| 39 | 772-786 | FYASLTYLRRGGGGS | MERCI | POSITIVE | 7 |
| 40 | 773-787 | YASLTYLRRGGGGSF | MERCI | POSITIVE | 7 |
| 41 | 774-788 | ASLTYLRRGGGGSFV | MERCI | POSITIVE | 7 |
| 42 | 775-789 | SLTYLRRGGGGSFVI | MERCI | POSITIVE | 7 |
| 43 | 840-854 | STRGRKCCRRKKHHH | MERCI | POSITIVE | 1 |

**Supplementary table S10. Refinement models of CTL and HTL MEVs.** CTL and HTL MEVs models were refined by GalaxyWEB server and used for further studies. After refinement in particular Rama favored residues increased significantly.

| Galaxy Refinement for CTL MEV | | | | | | |
|---|---|---|---|---|---|---|
| Model | GDT-HA | RMSD | MolProbity | Clash score | Poor rotamers | Rama favored |
| Initial | 1.00 | 0.00 | 3.336 | 120.1 | 1.8 | 87.6 |
| MODEL 1 | 0.9596 | 0.385 | 2.673 | 29.9 | 1.8 | 90.8 |
| Galaxy Refinement for HTL MEV | | | | | | |
| Initial | 1.00 | 0.00 | 3.725 | 167.4 | 3.4 | 86.2 |
| MODEL 1 | 0.9463 | 0.419 | 2.811 | 38.4 | 1.6 | 88.5 |

**Supplementary table S11. B Cell discontineous epitopes of CTL & HTL MEVs.**
Discontinuous B Cell epitopes predicted by ElliPro (IEDB) from CTL & HTL MEVs.

| CTL MEV Discontinuous epitopes residues | | | |
|---|---|---|---|
| S.No | Residues | Number of residues | Score |
| 1 | G1, I2, G3, D4, P5, V6, T7, C8, L9, K10, S11, G12, A13, I14, C15, H16, P17, V18, F19, C20, P21, R22, R23, Y24, K25, Q26, I27, G28, T29, C30, G31, L32, P33, G34, T35, K36, C37, C38, K39, K40, P41, E42, A43, I94, Q95, V545, L546, S547, C548, L549, P550, K551, E552, S559, R561, G562, R563, R568, K569 | 59 | 0.747 |
| 2 | G246, Y288, G289, G290, G291, G292, S293, M294, P295, S296, D297, D298, F299, S300, N301, T302, F303, G304, G305, G306, G307, S308, G321, S322, I323, M324, K325, K326, S327, K329, Y331, G332, G333, N357, I359, Y360, G361, G362, G363, G364, S365, A366, E367, Y389, G390, G391, G392, G393, S394, I395, A396, T397, V398, G407, S408, L409, E410, T411, D412, D413, Y414, N415, G416, I417, Y418, G419, G420, G421, G422, S423, E424, T425, D426, D427, Y428, N429, G430, I431, Y432, G433, G434, G435, G436, S437, S438, Q439, N440, L441, L442, G448, G449, G450, S451, T452, S453, D454, L455, D456, V458, F460, Y461, G462, G463, G464, G465, S466, F467, P468, I469, S470, R471, F473, N474, M475, Y476, G477, G478, G479, G480, S481, R482, G508, G509, S510, K511, Y512, Q514, D516, Q517, F520 | 130 | 0.695 |
| 3 | F56, G57, G58, G59, G60, S61, A62, Q63, A96, G102, G103, G104, G105, G116, G117, G118, G119, S120, L121, A122, M123, D124, E125, G126, Y127, G132, G133, G134, S135, T136, V137, Y138, H139, S141, A142, V143, Y144, G145, G146, G147, G148, S149, A150, V151, Y152, N153, N154, E155, F156, Y157, Y158, G159, G160, G161, G162, S163, A164, E165, N166, G173, G174, G175, G176, S177, N178, Y179, M180, Y181, F187, G188, T198, M199, L200, E201, F202, G203, G204, G205, G206, S207, T208, P209, F210, V211, D212, S213, R214, A215, Y216, G217, G218, G219, G220, S221, G231, G232, G233, G234, S235, T255, D256, V257, V258, Y259, G260, G261, G262, G263, S264, L265, V266, S267, D268, A269, K270, M271, L272, S273, G275, G276, G277, G278, S279, V280, S281, D282, A283, K284, M285, L286, S287 | 131 | 0.682 |
| HTL MEV Discontinuous epitopes residues | | | |
| S.No | Residues | Number of residues | Score |
| 1 | G1, I2, G3, D4, P5, V6, T7, C8, L9, K10, S11, G12, A13, I14, C15, H16, P17, V18, F19, C20, P21, Y24, K25, Q26, I27, G28, T29, C30, G31, L32, P33, G34, T35, K36, C37, C38, K39, K40, P41, E42, A43, A44, A45, K46, V47, Q48, M49, T50, Y51, N52, W53, T54, Q55, L57, Q58, Y61, G83, G84, G85, S86, L87, A88, L89, S90, K91, Y92, S113, V118, F119, G120, P121, G122, G123, G124, G125, S126, L127, S128, K129, Y130, L131, S132, D133, L134, L135, N160, L161, G162, G163, G164, G165, S166, K167, Y168, L169, F195, G196, P197, N198, L199, Q200, D201, G202, G203, G204, G205, S206, A207, S208, F209, S210, F235, G236, D237, G262, G263, G264, G265, S266, N267, D268, G283, G284, G285, S286, D287, M499, L500, S501, G502, G503, G504, G505 | 133 | 0.745 |
| 2 | S71, G319, V320, F321, G322, G323, G324, G325, R331, N333, W334, I335, S336, A337, G338, V339, F340, L341, G342, G343, G344, G345, S346, I347, P348, R349, E350, F351, M352, I353, Y354, D355, D356, V357, F358, I359, D360, N361, G362, G363, G364, G365, S366, F367, M368, I369, Y370, D371, D372, V373, F374, I375, D376, N377, T378, G379, R380, I381, G382, G383, G384, G385, S386, L387, S388, S389, D390, Q391, V392, A393, E394, L395, Q400, E401, G402, G403, G404, G405, S406, S407, S408, L433, A434, A435, A436, V437, Q438, E439, T440, G442, G443, G444, G445, S446, D447, Q448, A450, E451, A454, Q457, E458, T459, S460, A461, G462, G463, G464, G465, S466, Q467, V468, A469, E470, L471, A472, A473, A474, V475, Q476, E477, T478, S479, A480, G481, G482, G483, G484, G485, S486, N487, N488, G489, N490, S514, D515, A516, K517, M518, G522, G523, G524, G525, S526, G527, D553, A554, K555, M556, S558, Y559, A560, P561, G562, G563, G564, G565, A573, K574, M575, L576, S577, Y578, A579, P580, E581, G582, G583, G584, G585, S586, N587, D589, F601, G602, G603, G604, G605, S606, I607, D608, N609, F616, R640, N647, I648, H649, L650, L651, A652, E653, F654, F655, S656, F657, F658, R659, T660, F661, G662, G663, G664, G665, S666, I667, H668, L669, L670, A671, F673, T679, F680, G681, G682, G683, G684, G702, G704, G705, S706, E707, L708, A709, S710, F711, L712, M713, D714, R715, R716, V717, I718, R721, G722, G723, G724, G725, S726, L727, A728, S729, F730, L731, D733, P739, A741, G742, G743, G744, G745, S786, F787, V788, I789, F790, A792, S793, L794, T795, Y796, L797, R798, R799, G800, I801, E802, A803, A804, A805, K806, G807, I808, I809, N810, T811, L812, Q813, K814, Y815, Y816, R820, G821, G822, A825, V826, L827, S828, C829, L830, P831, K832, E833, E834, Q835, I836, G837, K838, T841, R842, G843, K845, C846, C847, R848, R849, K850, K851, H852, H853, H854, H855, H856 | 311 | 0.687 |
| 3 | D273, D292, I294, N295, I297, S298, A299, G300, V301, G302, G303 | 11 | 0.624 |
| 4 | G105, S106, A107, L108, S109, K110, Y111, E420, T421, G422, G423, G424, G425, S426, S427, D428 | 16 | 0.598 |
| 5 | G139, P140, N141, G142, G143, G144, G145, S146, S147 | 9 | 0.558 |

**Supplementary figure S1.** Workflow chart.

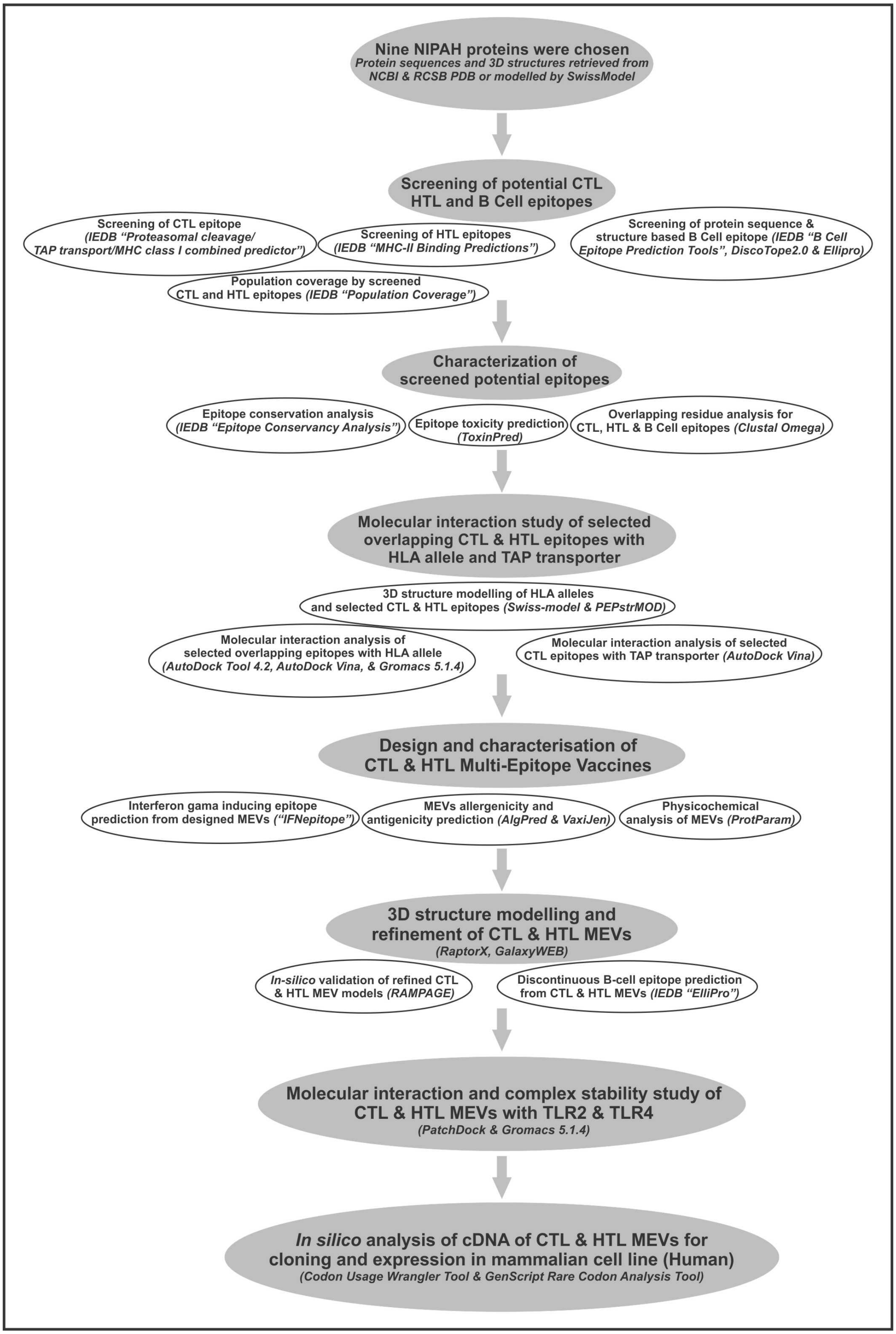

**Supplementary figure S2. (A)** Rg (radius of gyration) for the CTL epitope – HLA class I allele complexes, across the time window of 1 nano second. **(B)** Rg for the HTL epitope – HLA class II allele complexes, across the time window of 1 nano second.

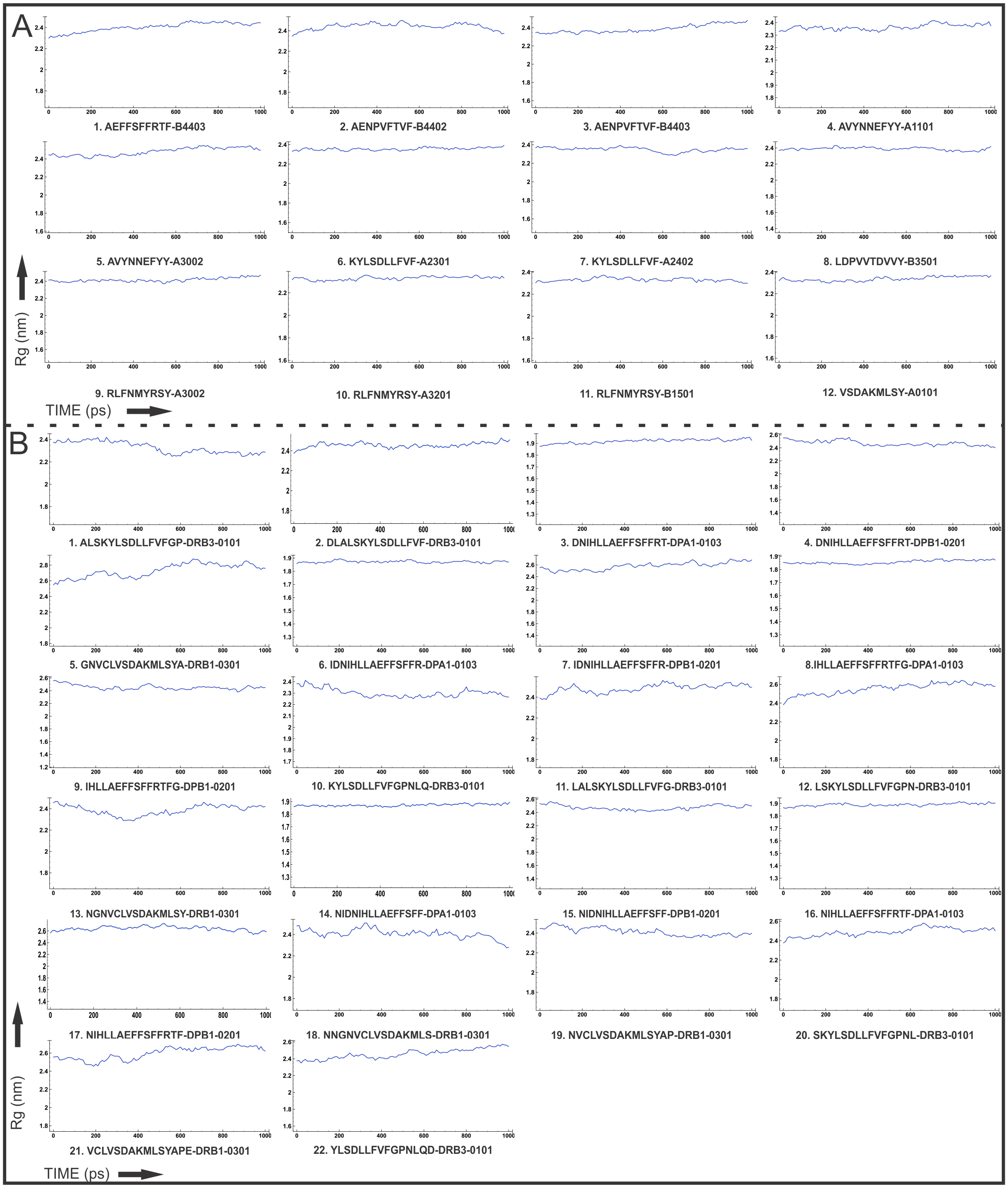

**Supplementary figure S3. (A)** RMS fluctuation in nanometers for all the atoms of the CTL epitope – HLA class I allele complexes. **(B)** RMS fluctuation in nanometers for all the atoms of the HTL epitope – HLA class II allele complexes.

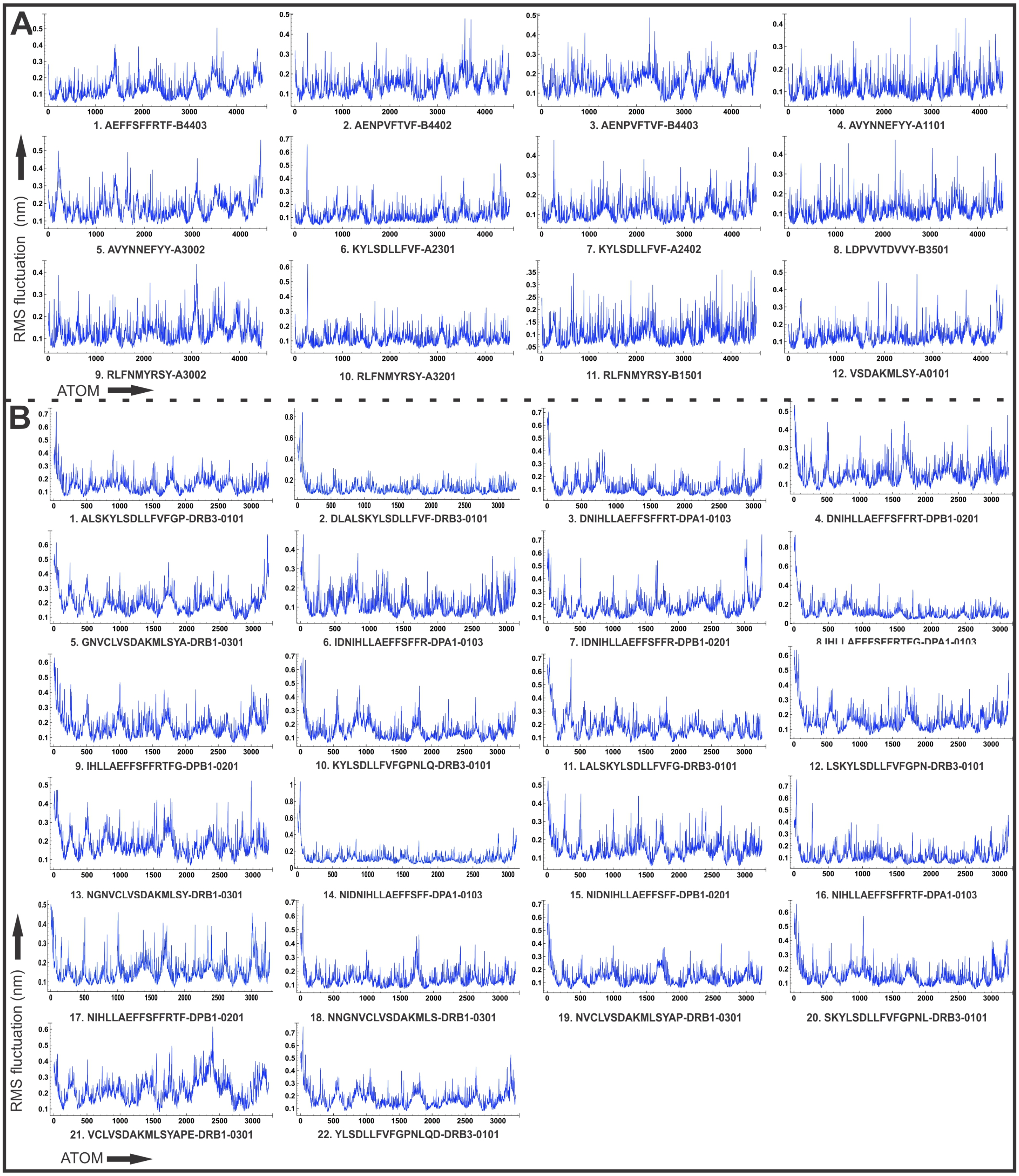

**Supplementary figure S4. (A)** B-Factor of CTL epitope – HLA class I allele complexes **(B)** B-Factor of HTL epitope – HLA class II allele complexes. Epitopes are shown in sticks and HLA alleles are shown in gray cartoons. B-factor is indicated by rainbow (VIBGYOR) colour, blue for stable region and red for most unstable region.

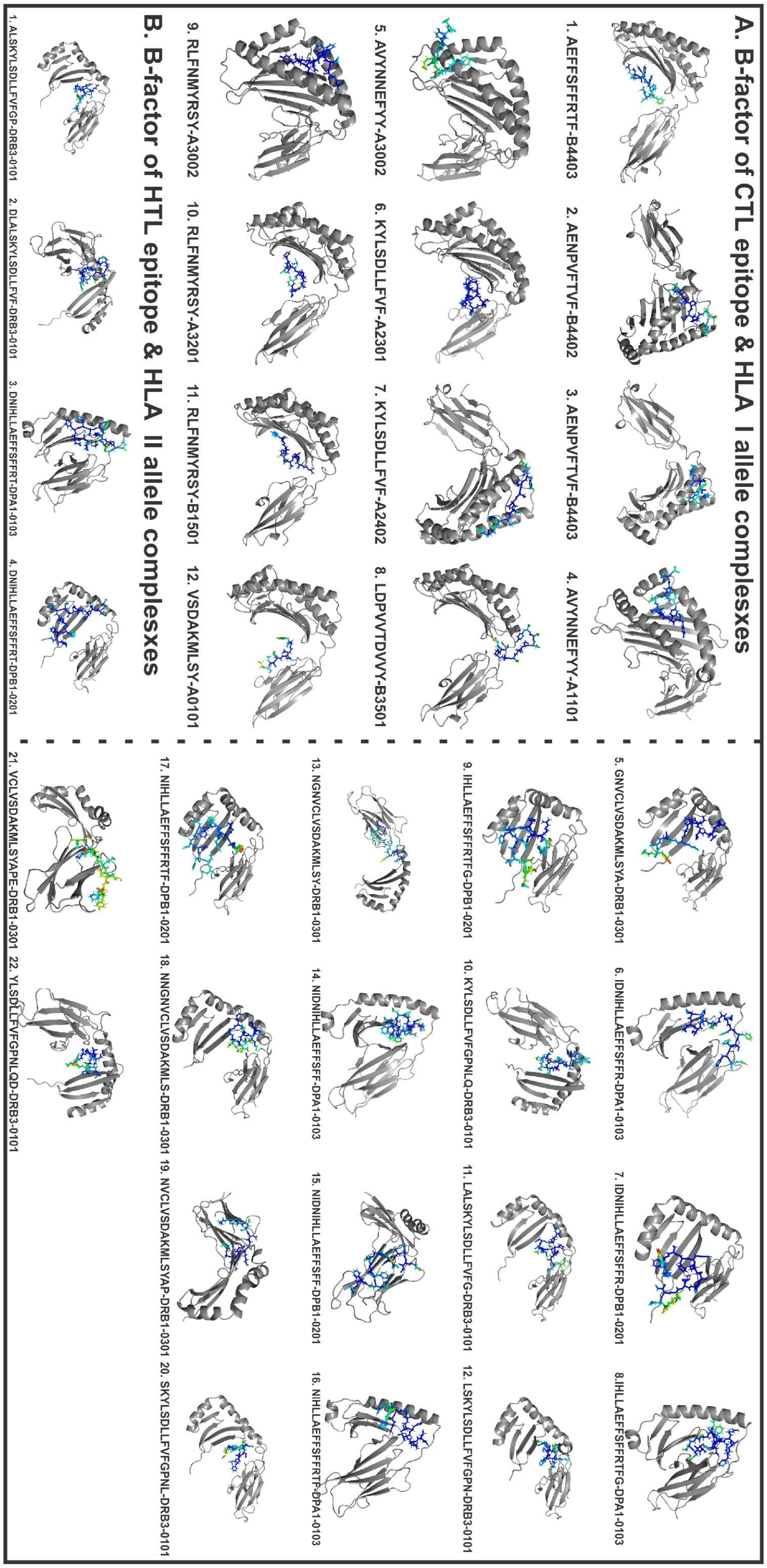
